# Amino Acid and Glucose Fermentation Maintain ATP Content in Mouse and Human Malignant Glioma Cells

**DOI:** 10.1101/2024.04.18.589922

**Authors:** Derek C. Lee, Linh Ta, Purna Mukherjee, Tomas Duraj, Marek Domin, Bennett Greenwood, Srada Karmacharya, Niven R. Narain, Michael Kiebish, Christos Chinopoulos, Thomas N. Seyfried

## Abstract

Energy is necessary for tumor cell viability and growth. Aerobic glucose-driven lactic acid fermentation is a common metabolic phenotype seen in most cancers including malignant gliomas. This metabolic phenotype is linked to abnormalities in mitochondrial structure and function. A luciferin-luciferase bioluminescence ATP assay was used to measure the influence of amino acids, glucose, and oxygen on ATP content and viability in mouse (VM-M3 and CT-2A) and human (U-87MG) glioma cells that differed in cell biology, genetic background, and species origin. Oxygen consumption was measured using the Resipher system. Extracellular lactate and succinate were measured as end products of the glycolysis and glutaminolysis pathways, respectively. The results showed that: 1) glutamine was a source of ATP content irrespective of oxygen. No other amino acid could replace glutamine in sustaining ATP content and viability; 2) ATP content persisted in the absence of glucose and under hypoxia, ruling out substantial contribution through either glycolysis or oxidative phosphorylation (OxPhos) under these conditions; 3) Mitochondrial complex IV inhibition showed that oxygen consumption was not an accurate measure for ATP production through OxPhos. The glutaminase inhibitor, 6-diazo-5-oxo-L-norleucine (DON), reduced ATP content and succinate export in cells grown in glutamine. The data suggests that mitochondrial substrate level phosphorylation in the glutamine-driven glutaminolysis pathway contributes to ATP content in these glioma cells. A new model is presented highlighting the synergistic interaction between the high-throughput glycolysis and glutaminolysis pathways that drive malignant glioma growth and maintain ATP content through the aerobic fermentation of both glucose and glutamine.

**Summary statement:** Malignant gliomas, regardless of cell of origin or species, rely on fermentation mechanisms for ATP production due to OxPhos insufficiency. Glucose and glutamine together are necessary and sufficient for dysregulated tumor cell growth, whereas OxPhos is neither necessary nor sufficient.

## Introduction

Gliomas are a common form of primary brain cancers which remain largely unmanageable. Malignant glioma is often diagnosed in older patients but can occur at any age with rising cases in children (Phillips et al., 2006; Tamimi and Juweid, 2017). Median survival is between 8-15 months with overall survival after 5 years remains below 10% (Omuro and DeAngelis, 2013; Fatehi et al., 2018; Seyfried et al., 2022). Patient outcome remains poor due in large part to the invasive characteristics in gliomas making complete surgical resection difficult (McCutcheon and Preul, 2021; Seyfried et al., 2022). Gliomas encompass numerous cell types that express glial, stem cell, and mesenchymal markers (Aum et al., 2014; Sidaway, 2017). Despite the extensive cellular and genetic heterogeneity, malignant glioma cells are more dependent on fermentation than on respiration for survival due the well-documented abnormalities in mitochondrial structure and function(Arismendi-Morillo, 2009; Seyfried et al., 2020).

Cancer cells exhibit a number of hallmarks including angiogenesis, apoptotic evasion, immune suppression, among others (Hanahan and Weinberg, 2000). Metabolic reprogramming was subsequently included as it became apparent that nearly all cancers exhibit similar aberrant metabolic phenotypes (Hanahan and Weinberg, 2011; Seyfried et al., 2020). Indeed, accumulating evidence shows that cancer is primarily a mitochondrial metabolic disease (Seyfried and Shelton, 2010; Coller, 2014; Seyfried et al., 2014; Wishart, 2015; Poljsak et al., 2019; Seyfried and Chinopoulos, 2021). Aerobic fermentation, later referred to as the Warburg effect, is the most well-known metabolic abnormality found in cancer cells (Warburg, 1956; Seyfried et al., 2014). Warburg first proposed a two-step process for the origin of cancer. The first step involved a chronic disruption of energy produced through oxidative respiration, while the second step involved a protracted replacement of insufficient respiratory energy with enhanced energy production through cytoplasmic fermentation (Burk and Schade, 1956; Warburg, 1956). Aerobic fermentation converts most glucose to lactate in tumor cells, regardless of oxygen availability, rather than fully oxidizing glucose through the TCA cycle. Healthy tissues perform this shunted utilization of glucose under hypoxic conditions or extreme energy demand such as high-intensity exercise (Hermansen and Stensvold, 1972) with the exception of certain populations of CNS cells, *e.g.* astrocytes which can metabolize glucose to lactate even in the presence of oxygen *in vivo* (Pellerin et al., 2007; Dobolyi et al., 2024). The question of whether tumor cells utilize oxidative phosphorylation (OxPhos), efficiently or otherwise, has been a continuous debate for nearly a century (Burk and Schade, 1956; Weinhouse, 1956; Aisenberg, 1961; Colowick, 1961; Kiebish et al., 2008; Hao et al., 2010; Marie and Shinjo, 2011). Recent studies show that all major cancers, including gliomas, have documented defects in the number, structure, and function of their mitochondria (Arismendi-Morillo and Castellano-Ramirez, 2008; Putignani et al., 2012; Deighton et al., 2014; Seyfried et al., 2020). These defects would compromise OxPhos function according to the foundational evolutionary principle that mitochondrial structure determines function (Lehninger, 1964; Cogliati et al., 2016; Miyazono et al., 2018; Baker et al., 2019; Glancy et al., 2020; Seyfried et al., 2020). A transition of energy production from respiration to fermentation would be essential for tumor cells to remain viable according to Warburg’s hypothesis.

In addition to glucose, glutamine is also recognized as important for the growth of many tumor cells including gliomas (DeBerardinis et al., 2007; Krall and Christofk, 2015; Bolzoni et al., 2016; Marquez et al., 2017). Glutaminolysis is considered an essential metabolic pathway for unbridled proliferation (McKeehan, 1982; Dang, 2010; Zhou et al., 2012; Oizel et al., 2017; Still and Yuneva, 2017; Nilsson et al., 2020). Glutamine is the most abundant amino acid in humans, ranging from 0.4-1.0 mM in the circulation (Betz and Goldstein, 1986; Smith and Wilmore, 1990). In contrast to serum and extra neural tissues, glutamine is tightly regulated in the brain through its involvement in the glutamate–glutamine cycle of neurotransmission (McKenna, 2007; Schousboe et al., 2014; Sonnewald and Schousboe, 2016). Radiation-induced disruption of this cycle can provide neoplastic glioma cells increased access to glutamine (Seyfried et al., 2019). Besides serving as a metabolic fuel for the neoplastic cells, glutamine is also an important fuel for cells of myeloid lineage, which include microglia, and especially the highly invasive mesenchymal cells in glioma (Huysentruyt et al., 2011). The regulation of glutamine metabolism in the tumor microenvironment is therefore important for managing the tumor (Seyfried et al., 2022).

In this report, we used a luciferin-luciferase bioluminescence ATP assay to measure the influence of amino acids and glucose on the growth and survival of mouse and human glioma cell lines. We tested the relationship between oxygen consumption and ATP content in two mouse glioma cell lines, VM-M3 and CT-2A, and in the human glioma cell line, U-87MG. The three lines differ from each other in cell biology, genetic background, and in species of origin, but are similar to each other in having mitochondrial defects that would compromise OxPhos efficiency (Kiebish et al., 2008; Sun et al., 2019). Our results show that in addition to glucose, glutamine can also be fermented in the presence or absence of oxygen for ATP production through the glutaminolysis pathway in these three malignant glioma cell lines.

## Materials and Methods

### Cell lines

The VM-M3 tumor arose spontaneously in the cerebrum of an adult mouse of the VM/dk inbred strain maintained in the Biology Department at Boston College. A cell line was established from the VM-M3 tumor as previously described (Huysentruyt et al., 2008). The VM-M3 cell line manifests most of the growth characteristics typical of human glioma including distal brain invasion using the secondary structures of Scherer (Shelton et al., 2010). The highly angiogenic CT-2A tumor was produced from 20-methancolanthrene in the cerebral cortex of a C57BL/6J mouse as previously described (Seyfried et al., 1992). CT-2A has several characteristics in common with the neoplastic neural stem cells found in human glioma (Binello et al., 2012). CT-2A has been described with some of the characteristics of a highly malignant anaplastic astrocytoma (Seyfried et al., 1992). U-87MG was isolated from tumor tissue from a 44-year-old female glioma patient in 1966 and serves as a common model for human glioma (Ponten and Macintyre, 1968). The VM-M3, CT-2A, and U-87MG cell lines were transduced with a lentivirus vector containing the firefly luciferase gene under the control of the cytomegalovirus promoter (e.g., VM-M3/Fluc) as previously described (Sena-Esteves et al., 2004; Huysentruyt et al., 2008). All cell lines are mycoplasma free.

### Culture conditions

Stock plates of tumor cells were cultured in DMEM (D5796; Sigma-Aldrich, St. Louis, MO) supplemented with 10% fetal bovine serum (FBS; FS-0500-AD, lot: S12E22AD1; Atlas Biologicals, Fort Collins, CO) containing 25.0 mM glucose, 4.0 mM glutamine, and 1% penicillin-streptomycin at 37°C. Experimental media was made from powdered DMEM (D5030; Sigma-Aldrich). Glucose and glutamine were added for each experiment. PBS solutions (21600-044; Gibco) were fortified with 0.37 g/L sodium bicarbonate. Potassium cyanide, sodium azide, and 6-diazo-5-oxo-L-norleucine (Sigma-Aldrich) were stored at 4°C. In some of the experiments, cells were grown in a sealed chamber (Biospherix, Parish, NY) equilibrated with N_2_ under normoxia (21% O2) or hypoxia (0.1% O_2_). The 20 amino acids used are listed in *Supplemental Table 1*.

### Luciferin-luciferase Bioluminescence Assay

Tumor cells were seeded and allowed to settle overnight in seeding media for up to 18 hours. The following morning, seeding media was removed and replaced by various experimental media. Luciferin (10 mg/mL, Syd Labs, Hopkinton, MA) was added to each well containing 5.0 × 10^3^ luciferase-tagged cells and allowed to incubate at room temperature for 1 minute as previously described (Huysentruyt et al., 2008). Plates (96-well) were read with the Ami HT (Spectral Instruments, Tucson, AZ) device after a five-minute exposure. Aura software version 2.3.1 was used for quantification and visualization of bioluminescence. Photon values were obtained by ROI selection of each well as per manufacturer’s instructions using Aura software.

### Trypan Blue Exclusion Assay

Trypan blue exclusion was used to quantify tumor cell viability (Crowley et al., 2016). The tumor cells were seeded as above. After incubation in the experimental media, a 0.4% trypan blue solution (10 µL) was added to each well. After 10 minutes of exposure, the wells were washed twice with sterile PBS at room temperature. The number of trypan blue positive and negative cells was evaluated with a brightfield 20 X objective lens using an EVOS microscope (Life Technologies, Carlsbad, CA).

### Calcein-AM and Ethidium Homodimer III Assay

Calcein-AM and ethidium homodimer III (EthD-III) were used to quantify tumor cell viability (Decherchi et al., 1997). The tumor cells were seeded as above. After 24 hours of incubation in experimental media, a 100 µL solution containing 4 mM calcein-AM and 2 mM EthD-III (100 µL) was added to each well. After 30 minutes of incubation at 37°C, cells were imaged using a 20 X objective lens with either a Texas Red or GFP filter cube on an EVOS microscope (Life Technologies, Carlsbad, CA). The number of calcein-AM positive and EthD-III negative cells were counted to determine the percentage of viable cells.

### Succinate Assay

Extracellular succinic acid levels were measured using a colorimetric assay (BioAssay Systems, Hayward, CA) as per manufacturer’s instructions. Cells were seeded as described above. The linear detection range of the assay was 10 to 400 µM. Aliquots (20 µL) were taken after 6 hours. The absorbance was read at 570 nm after a 30-minute incubation period at room temperature. Concentrations were calculated using absorbances following manufacturer’s recommendations. Succinic acid was also measured using LC-MS as described below.

### Lactate Assay

Extracellular lactic acid levels were measured using a colorimetric assay (Eton Biosciences, San Diego, CA) as per manufacturer’s instructions. Cells were seeded as described above. Aliquots (50 µL) of media were moved to a new plate and mixed with 50 µL of L-Lactate Assay Solution. Absorbance was read at 490 nm after a 30-minute incubation period at room temperature. Absorbance values were compared to those on a standard curve to determine lactate concentration.

### Oxygen Consumption Assay

Oxygen consumption was measured continuously using the Resipher instrument (Lucid Scientific, Atlanta, GA) in 96-well Falcon flat-bottom plates. Cells were seeded at a density of 2.0 × 10^5^ cells/well. The sensing lid and Resipher device were placed on top of the 96-well plate for five minutes prior to the start of the experiment. Experimental media containing glutamine with sodium azide or potassium cyanide was added. The sensing lid was placed on top of the Falcon plate and put into the HEPA 100 incubator (Thermo Fisher Scientific, Waltham, MA) maintained at 5% CO_2_ and 88% relative humidity. Oxygen consumption rate (OCR) was monitored in the incubator for up to four days. Data was analyzed using the Resipher web application.

### Complex IV Activity Assay

Protocol for determining complex IV activity was adapted from Doczi *et al.,* (Doczi et al., 2023). Briefly, 2.0 × 10^4^ cells were seeded in clear bottom 96-well plates. Culture medium was switched with a solution containing: 20 mM KH_2_PO_4_, 0.1 µM myxothiazol, 0.45 mM n-dodecyl-β-maltoside, pH 7.0) with or without 1 mM KCN. Reaction was started with 50 µM ferrocytochrome C after 10 minutes of temperature equilibration at 37°C. Cytochrome C was reduced on the day of measurement using dithionite. Absorbance was recorded at 550 nm.

### Liquid Chromatography-Mass Spectrometry (LC/MS)

LC/MS (Agilent 1200 series liquid chromatography system) was used for analysis of labeled and unlabeled metabolites. Extracellular media was collected from VM-M3 cells that were cultured for 17 hours in media containing 1 mM C^13^-glutamine and 1 mM glutamine. Samples were flash frozen using dry ice until processing. Cold milliQ-purified water (150 µL) and 1.0 mL of methanol (Fischer Scientific Optima LC/MS, Waltham, MA). After brief vortexing, the combined solution was centrifuged at *10,000 x g* for 10 minutes. The supernatant was dried using a SpeedVac (Eppendorf, Hamburg, Germany) at 30°C. The media extract was reconstituted in 100 µL of methanol and filtered prior to sample loading for LC/MS analysis using a triple-quad method (Agilent 6460). The SGE Analytical Science ProteCol HPLC C18 Q103 150 mm x 2.1 mm ID 3 um 100A column used during the extraction procedure. Formic acid (0.2%) was used for Mobile Phase A. The following parameters were used: Gas temperature, 320°C; Gas flow, 10 L/minute; Nebulizer, 50 PSI; Sheath gas temperature, 315°C; Sheath gas flow, 11 L/minute; Capillary voltage, 4000 volts.

### Metabolite Extraction and Metabolomics Quantitative Analysis

Metabolite extraction using media samples was achieved using cold 80% methanol stored at −20°C. We used 10 mL of cell media for metabolite extraction which was extracted three times with cold 80% methanol (500 mL, 100 mL, and 50 mL to achieve dilutions of 50x, 10x, and 5x, respectively). This methodology allowed enhanced detection of low abundance target metabolites to achieve calculated concentrations within the linear calibration ranges of the instrument. All samples were stored at −20°C for 1 hour during full metabolite extraction. Supernatant extracts were transferred for analysis via multiple reaction monitoring (MRM) hydrophilic interaction chromatography-liquid chromatography-tandem mass spectrometry (HILIC-LC-MS/MS). Separation was performed using a NEXERA XR UPLC system (Shimadzu, Columbia, MD) coupled with the Triple Quad 5500 System (AB Sciex, Framingham, MA).

Quality control was performed using a metabolite standards mixture and pooled samples approach. A standard quality control sample containing a mixture of amino and organic acids was injected daily to monitor mass spectrometer calibration. A pooled quality control sample was obtained by taking an aliquot of all samples injected daily to determine the optimal dilution of the batch samples to validate metabolite identification and peak integration. Quantitative analysis was performed using a set of mixed metabolite standards at known concentrations analyzed along with the set of media samples. As previously stated, standard quality control samples containing a mixture of amino and organic acids was injected daily to monitor mass spectrometer response. Linear regression was applied, and the concentration of each target metabolite was calculated in the original cell media.

### Global Metabolomics Data Collection and Applied Enrichment and Pathway Analyses

Global metabolomics was conducted to analyze the entirety of extracellular metabolites from various media conditions. The global metabolomics analysis was applied to extracts from the 50x dilution sample set as described above. Supernatant extracts were split, and analyses were performed using gas chromatography-mass spectrometry (GC-MS), reversed-phase liquid chromatography-mass spectrometry (RP-LC-MS), and HILIC-LC-MS/MS with applied multiple reaction monitoring. RP-LC-MS separation was performed using the NEXERA XR UPLC system (Shimadzu, Columbia, MD) coupled with a Triple TOF 6600 System (AB Sciex, Framingham, MA). GC-MS aliquot separation was performed using an Agilent 7890B gas chromatograph (Agilent, Palo Alto, CA) interfaced to a Time-of-Flight Pegasus HT Mass Spectrometer (Leco, St. Joseph, MI). The GC system was fitted with a Gerstel temperature-programmed injector cooled injection system (model CIS 4). An automated liner exchange (ALEX) (Gerstel, Muhlheim an der Ruhr, Germany) was used to eliminate cross-contamination from the sample matrix occurring between runs. A standard quality control sample containing a mixture of amino and organic acids were injected daily to monitor mass spectrometer response. A pooled quality control sample was obtained by taking an aliquot of all samples for daily injection to determine the optimal dilution of the batch samples and validate metabolite identification and peak integration. Raw data were manually inspected, merged, inputted, and normalized by the sample median. Global metabolomic data were analyzed as previously described (Tolstikov et al., 2014; Chi et al., 2022). Identified metabolites were subjected to pathway analysis with MetaboAnalyst 4.0, using the MSEA module consisting of an enrichment analysis that relies on measured levels of metabolites and pathway topology. MetaboAnalyst 4.0 provides visualization (heatmap) of the identified metabolic pathways. Accession numbers of detected metabolites (HMDB, PubChem, and KEGG Identifiers) were generated, manually inspected, and utilized to map the canonical pathways.

### Western Blot Analysis of Pyruvate Kinase Isoform Protein Expression

Western blots were performed to determine the presence of PKM1 and PKM2 as described previously (Mukherjee et al., 2019). Briefly, VM-M3 Cells were homogenized in ice-cold lysis buffer containing 20 mM Tris-HCl (pH 7.5), 150 mM NaCl, 1 mM Na_2_EDTA, 1 mM EGTA, 1% Triton, 2.5 mM NaPPi, 1 mM a-glycerophosphate, 1 mM Na_3_PO_4_, 1 Ag/mL leupeptin, and 1 mM phenylmethylsufonyl fluoride. Lysates were transferred to 1.5 mL Eppendorf tubes, mixed on a rocker for 1 hour at 4°C and then centrifuged at *8000 x g* for 20 minutes. Supernatants were collected and protein concentrations estimated using the Bio-Rad detergent-compatible protein assay. Total protein (100 ug) from each lysate was denatured with SDS-PAGE sample buffer (63 mM Tris-HCl (pH 6.8), 10% glycerol, 2% SDS, 0.0025% bromophenol blue, and 5% 2-mercaptoethanol). Sample was resolved by SDS-PAGE on 4 to 12% Bis-Tris gels (Invitrogen, Waltham, MA). Proteins were transferred to a polyvinylidene difluoride immobilon TM-P membrane (Millipore, Burlington, MA) overnight at 4°C and blocked in 5% nonfat powdered milk in TBS with Tween 20 (pH 7.6) for 1 hour at room temperature. Membranes were probed with PKM1 or PKM2 primary antibodies (Cell Signaling Technology, Danvers, MA) overnight at 4°C with gentle shaking. The blots were then incubated with the anti-rabbit secondary antibody (Cell Signaling Technology, Danvers, MA) for 1 hour at room temperature. Bands were visualized with enhanced chemiluminescence.

### Statistics

Statistical analyses and data plotting were performed using GraphPad Prism 9. Data were analyzed by one-way ANOVAs with Tukey’s, Sidak’s, and Dunnett’s multiple comparison post-hoc analysis, or unpaired student’s t-tests. Pearson’s correlation tests were performed to assess the relationship between cell number and bioluminescence. Pearson’s correlation coefficient is reported as R^2^ with the 95% confidence interval. Statistical significance was considered when p ≤ 0.05 and all data are shown as mean ± SEM. Asterisks indicate the following: * p ≤ 0.05; ** p ≤ 0.01,; *** p ≤ 0.001; and **** p ≤ 0.0001 unless otherwise stated.

## Results

The aim of this research was to determine, 1) if amino acid fermentation could contribute to ATP content in mouse and human glioma cells and, 2) if oxygen consumption was an accurate marker for ATP production through OxPhos in these cells. A luciferin-luciferase bioluminescence ATP assay was used to measure the influence of amino acids, glucose, and oxygen on ATP content, viability, and proliferation in mouse (VM-M3 and CT-2A) and human (U-87MG) glioma cells. The luciferin-luciferase system was used to measure ATP content *in vitro* as previously described (Manfredi et al., 2002). Luciferin will react with luciferase to produce oxyluciferin and light in a viable cell. The amount of light is based on the amount of ATP present in the sample (**Figure S1A**). A significant linear relationship was observed (R^2^ = 0.9786) between the number of VM-M3 luciferase-tagged cells and the emitted bioluminescence indicating that bioluminescence is an accurate marker for cell number and viability (**Figure S1B**).

### Glucose and glutamine are both required for sustained viability and proliferation of VM-M3 cells

VM-M3 cells were stably transfected with the luciferase reporter (*see Methods).* To evaluate the influence of glucose and glutamine on VM-M3 cell viability, the cells were cultured in serum-free DMEM containing either 12.0 mM glucose and 2.0 mM glutamine (positive control), glucose alone, glutamine alone, or in basal media (DMEM alone; negative control). Bioluminescence increased from the initial seeding value over 48 hours in media containing both glucose and glutamine (**Figure 1A**). Bioluminescence increased at 24 hours in VM-M3 cells cultured in media containing glutamine alone but decreased by 48 hours. In contrast to media containing glutamine alone, bioluminescence continually decreased in media containing glucose alone over 48 hours like that seen in the negative control. Similar results were obtained after 24 hours using trypan blue exclusion and calcein-AM/ethidium homodimer-III staining assays under identical media conditions in VM-M3 and CT-2A cells (**Figure S2A-D**). Viability of U-87MG cells did not significant change after 24 hours (**Figure SE-F**), likely due to differences in basal metabolic rate between mouse and human cells (Terpstra, 2001). These findings show that glucose and glutamine are required to maintain robust viability and proliferation over 48 hours. To further evaluate the effect of glucose and glutamine on VM-M3 cell bioluminescence, the VM-M3 cells were incubated in varying concentrations of glucose and glutamine (**Figure 1B**). Bioluminescence was measured after incubation in media containing glutamine concentrations of 2.0, 1.5, 1.0, 0.5, 0.1, and 0.0 mM while maintaining glucose concentration constant for 24 hours. No significant reduction in bioluminescence was observed over glutamine concentrations from 2.0 to 0.5 mM. Bioluminescence was significantly reduced when glutamine was lowered to 0.1 mM. A larger reduction in bioluminescence was observed in the absence of glutamine. Bioluminescence was again measured after incubation in medias containing glucose at concentrations of 12.0, 6.0, 3.0, 1.0, 0.5, or 0.0 mM while maintaining glutamine concentration constant for 24 hours (**Figure 1B**). Bioluminescence reduced significantly when the glucose concentration was lowered from 12.0 mM to 1.0 mM. These findings, viewed collectively, suggest that glutamine is a more essential metabolite than glucose for maintaining bioluminescence and viability of VM-M3 cells over 24 hours.

**Figure 1.**
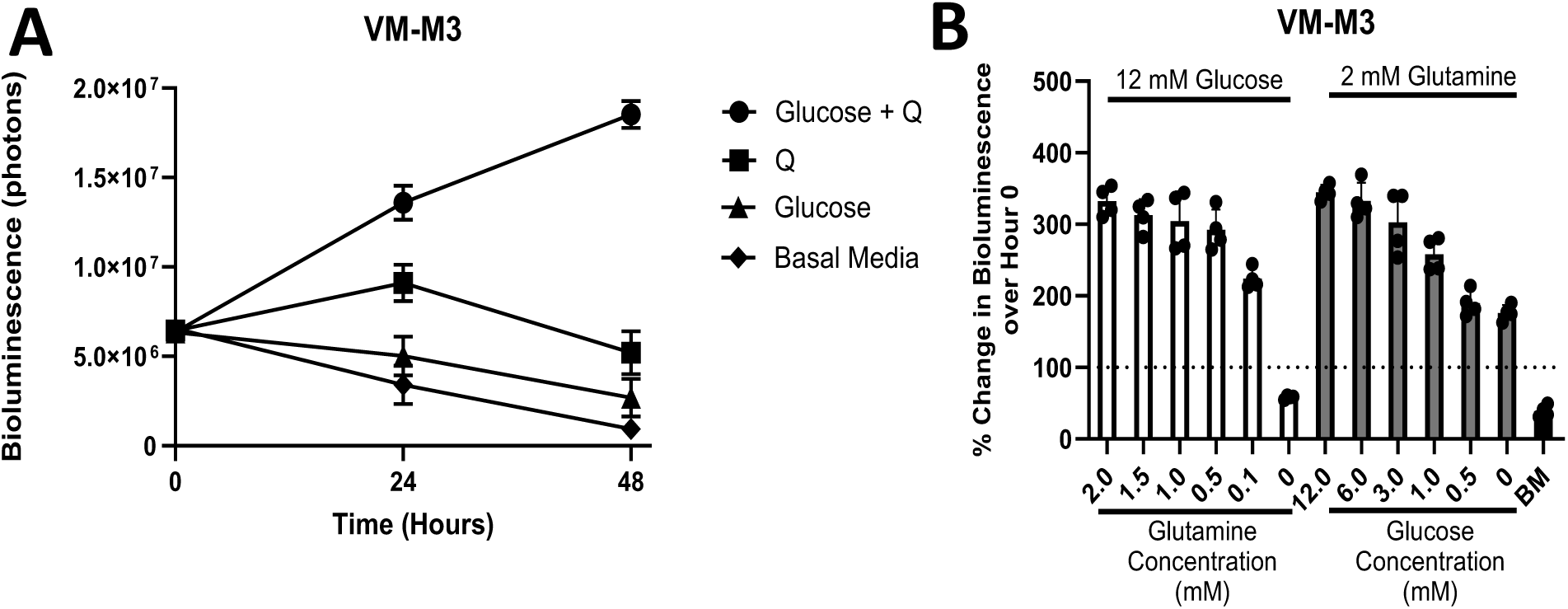
Influence of glucose and glutamine on the bioluminescence of VM-M3 cells. A) Cells were seeded at a density of 5.0 × 10^3^ and cultured for 48 hours with indicated media compositions. Glucose was added at 12 mM and glutamine (Q) added at 2 mM. Basal media represents DMEM with no added glucose or glutamine. Data are shown as mean ± SEM with an n=3 independent experiments for each time point. B) Bioluminescence of cells cultured in varying glucose and glutamine concentrations after 24 hours. Data are shown as mean percent increase in cell number relative to the hour 0 time point. BM indicates basal media (DMEM without added glucose or glutamine). Bars represent means of 3 independent experiments of 4 technical replicates per group.

### Lactate production and pyruvate kinase expression in VM-M3 cells

In contrast to the pyruvate kinase isoform (PKM1), where ATP content is linked to lactate production in the glycolytic pathway, the pyruvate kinase 2 isoform (PKM2) is a less active isoform in which lactate is produced independent of ATP content (Liu and Vander Heiden, 2015; Chinopoulos, 2020). Pyruvate kinase muscle isozymes 1 and 2 (PKM1 and PKM2) were examined by western blot analysis. VM-M3, CT-2A, and U-87MG cells expressed both PKM1 and PKM2 isoforms (**Figure S3A**). The majority of extracellular lactate was observed in the presence of glucose but not in the presence of glutamine in VM-M3, CT-2A, and U-87MG cells (**Figure S3B-D**). These findings indicate that less lactate production is derived from glutamine than from glucose and that lactate production may not be an accurate marker for ATP production through glycolysis in cancer cells that express both PKM1 and PKM2 isoforms.

### VM-M3 cells depend more on glutamine than on other amino acids for ATP content

To determine which amino acids are most critical for the survival of VM-M3 cells, we measured bioluminescence in cells that were individually supplemented with each amino acid over 48 hours. Several amino acids were tested at concentrations of 1.0, 2.0, and 4.0 mM. No differences in bioluminescence were found between these concentrations in preliminary experiments (data not shown). Amino acids were added to basal media (DMEM 5030) at 4.0 mM to ensure that the availability of each amino acid was not a limiting factor for ATP content for up to 48 hours. The basal media contained most amino acids except glutamine, glutamate, alanine, aspartate, asparagine, and proline. Bioluminescence of VM-M3 cells decreased by 64% after 24 hours in basal media indicating that the amino acids present in the basal media were unable to maintain the viability of this cell line (**Figure 2A**). Glutamine (+ 91%), glutamate (+ 30%), and lysine (+ 10%) were the only amino acids that increased bioluminescence over baseline after 24 hours, whereas glutamine was the only amino acid that could maintain bioluminescence for over 48 hours. Similar findings were obtained for the CT-2A and U-87MG cell lines among a smaller set of amino acids (**Figure S4A-B**). Methionine (− 89%) or valine (− 88%) significantly reduced bioluminescence relative to basal media alone after 24 hours. A full quantitative overview of the influence of each amino acid evaluated is shown in *Table 1*.

**Figure 2.**
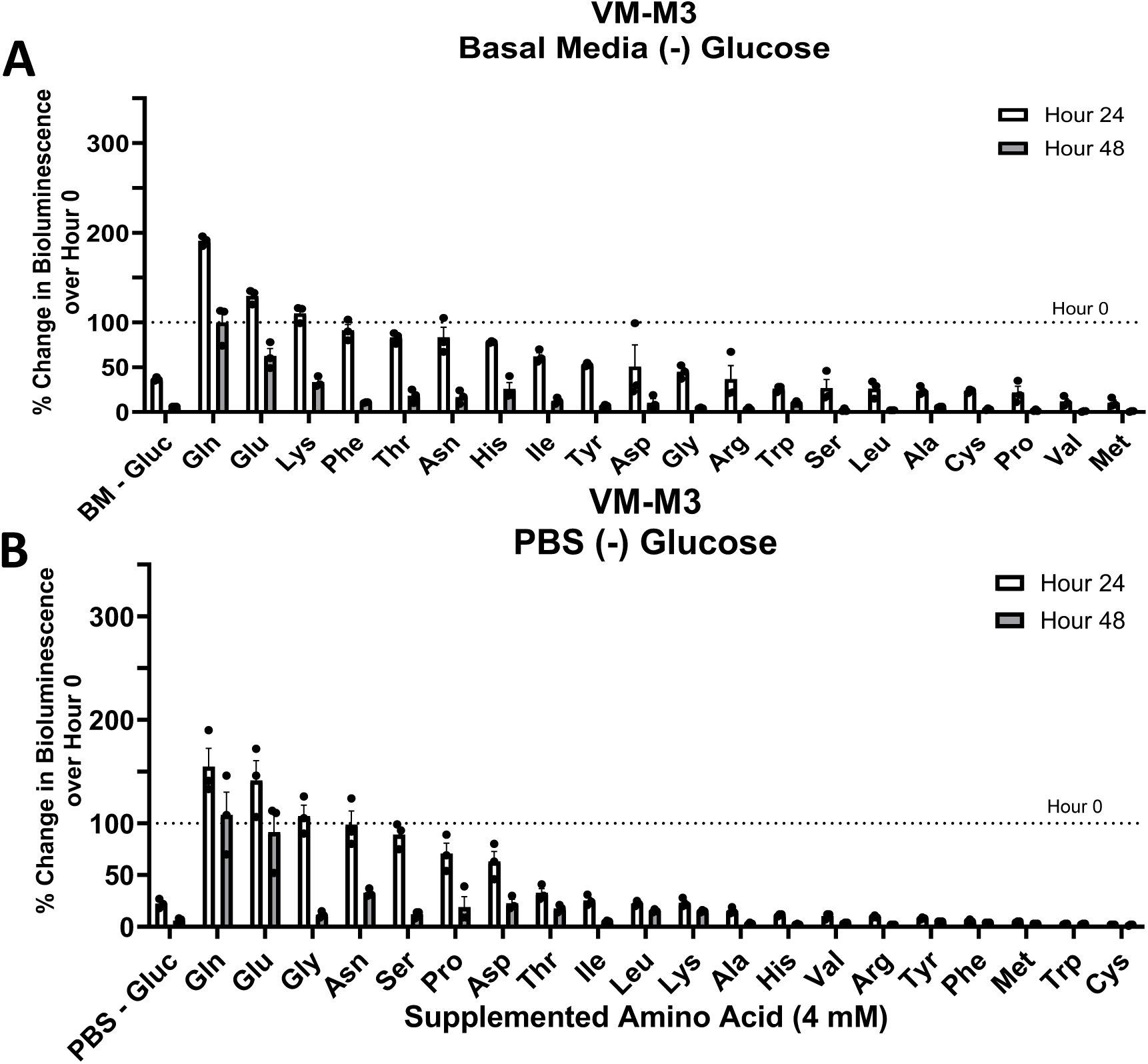
Influence of supplemented amino acids in basal media or phosphate buffered saline. A) VM-M3 cells cultured in basal media or B) phosphate buffered saline (PBS) with the indicated amino acid supplemented at 4 mM. The bioluminescence value at hour 0 is represented by a dashed line. Amino acids are ordered by descending percent change at hour 24. Cells were seeded at a density of 5.0 × 10^3^ cells/well. Quantitative values and statistical analysis can be found in *Table 1* (2A) and *Table 2* (2B). Values are shown as mean ± SEM with three independent experiments.

Next, individual amino acids were supplemented in phosphate buffered saline (PBS) to examine the specific contribution of each amino acid to bioluminescence under the most restrictive growth condition. Although PBS is an artificial environment, containing no metabolites, this system is highly effective in isolating the contribution of individual amino acids. A 78% decrease in bioluminescence was seen after 24 hours when VM-M3 cells were cultured in PBS alone (**Figure 2B**; *Table 2*). The addition of glutamine (+ 55%), glutamate (+ 42%), or glycine (+ 7%) increased bioluminescence, relative to the baseline, after 24 hours. Glutamine was the only amino acid that increased bioluminescence above baseline after 48 hours. On the other hand, methionine (− 96%), tryptophan (− 97%), or cysteine (− 98%) demonstrated significant inhibitory effects compared to PBS alone. These findings indicate that glutamine, compared to the other common amino acids tested, was most effective in sustaining bioluminescence in VM-M3 cells

### Glucose and glutamine synergize for the long-term proliferation of glioma cells

The proliferation of VM-M3 cells in basal media was significantly greater in the presence of glucose and glutamine than in the presence of either metabolite alone demonstrating a robust synergistic interaction between glucose and glutamine (**Figure 1A**). We sought to test whether glucose exhibited similar synergistic effects with other amino acids. In this aim, we evaluated the influence of individual amino acids (supplemented at 4.0 mM) in basal media containing 12.0 mM glucose over 48 hours. Besides glutamine (+ 119%), only lysine (+ 64%), isoleucine (+ 13%), and glutamate (+ 4%) increased bioluminescence over 24 hours (**Figure 3A**; *Table 3*). Glucose and glutamine were the only amino acid combination that could maintain bioluminescence above baseline after 48 hours.

**Figure 3.**
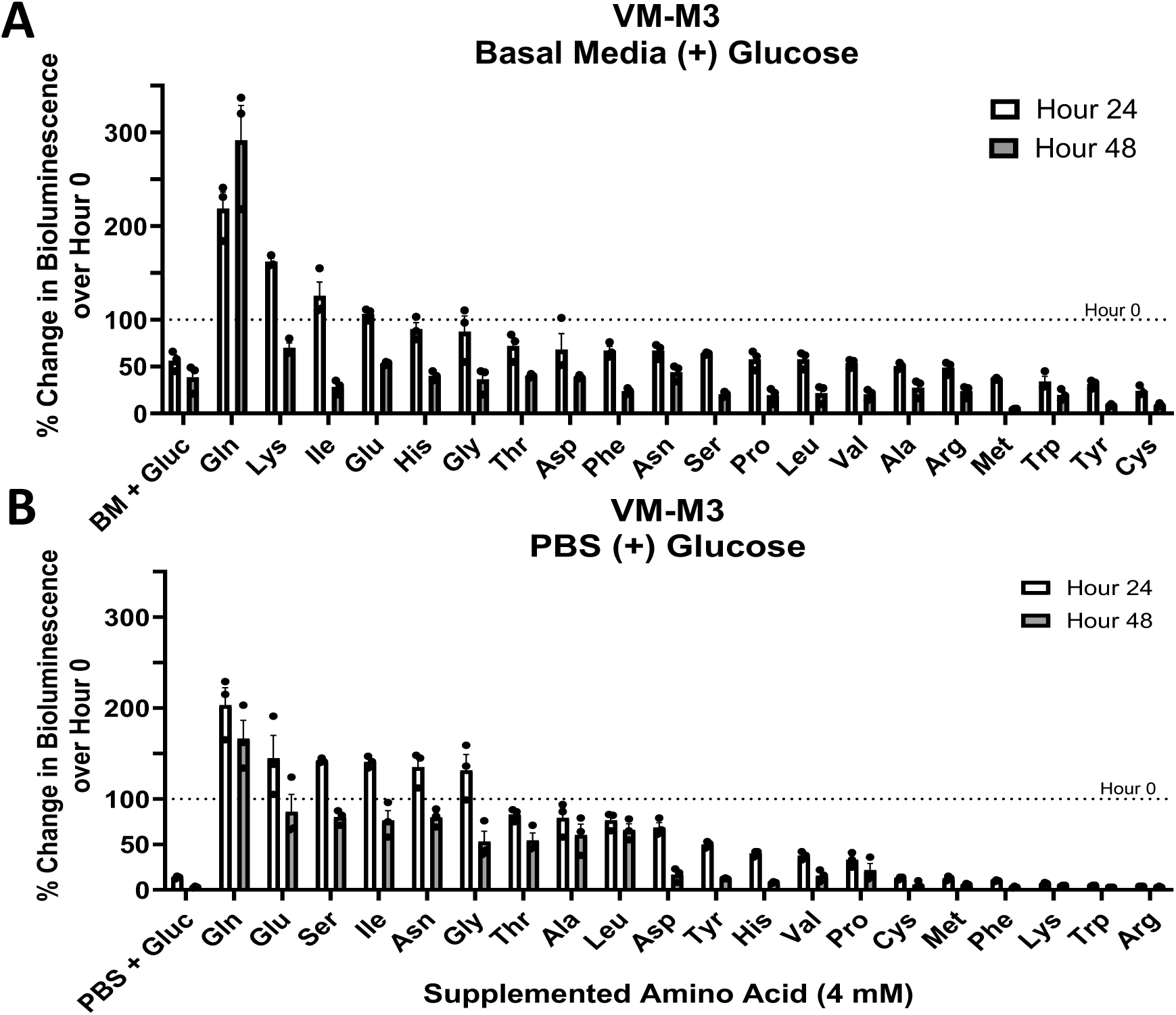
Influence of supplemented amino acids with glucose in basal media or phosphate buffered saline. A) VM-M3 cells cultured in basal media or B) PBS in 12mM glucose with the indicated amino acid supplemented at 4mM. The bioluminescence value at hour 0 is represented by a dashed line. Amino acids are ordered by descending percent change at hour 24. Cells were seeded at a density of 5.0 × 10^3^ cells/well. Quantitative values and statistical analysis can be found in *Table 3* (3A) and *Table 4* (3B). Values are shown as mean ± SEM with three independent experiments.

Next, VM-M3 cells were cultured in PBS containing 12.0 mM glucose to assess the synergy between amino acids and glucose alone (**Figure 3B**; *Table 4*). Amino acids that stimulated bioluminescence above baseline after 24 hours were: glutamine (+ 103%), glutamate (+ 45%), serine (+ 42%), isoleucine (+ 41%), asparagine (+ 35%), or glycine (+ 31%). As in basal media, glutamine (+ 66%) was the only amino acid that synergized with glucose to remain above baseline after 48 hours in PBS. The lower bioluminescence observed between glucose and glutamine in PBS versus basal media after 48 hours is likely attributable to the contribution of the additional co-factors such as amino acids and micronutrients that are also present within the basal media. These results suggest that glutamine is required for the sustained proliferation of VM-M3 cells over 24 hours.

Cysteine and methionine, two sulfur-containing amino acids, were consistently inhibitory across multiple conditions in VM-M3 cells (**Figures 2A-B, 3A-B**). This observation is inconsistent with previous studies suggesting that methionine-restriction (Chaturvedi et al., 2018) may be inhibitory for cancers through epigenetic mechanisms (Wanders et al., 2020). Although not explored in the present study, it is possible that the observed toxicity of high concentrations of cysteine and methionine may be attributed to the sulfur present in these amino acids. Further studies would be needed to test this hypothesis.

### Oxygen consumption rate (OCR) is not an accurate measure for ATP content through OxPhos in VM-M3, CT-2A, and U-87MG cells

To determine if oxygen consumption rate (OCR) is an accurate measure of ATP content in the absence of glucose, we cultured the VM-M3, CT-2A, and U-87MG cells with glutamine in the presence of sodium azide (Az), a complex IV inhibitor. CT-2A and U-87MG cells were used to investigate the effect of mitochondrial inhibitors in gliomas that differ in genetic background and species-of-origin. OCR and bioluminescence measurements were conducted in tandem. Two concentrations of sodium azide, 200 µM, and 500 µM, were used to account for potential cell-specific differences in response to complex IV inhibition. Mean oxygen consumption was reduced by 33% and 59%, respectively, in the VM-M3 cells (**Figure 4A**). Bioluminescence was measured simultaneously with no significant changes in the presence of 200 µM or 500 µM azide. These findings indicate that the reduction in ATP content in these cells was not proportional to the reduction in oxygen consumption under these conditions.

**Figure 4.**
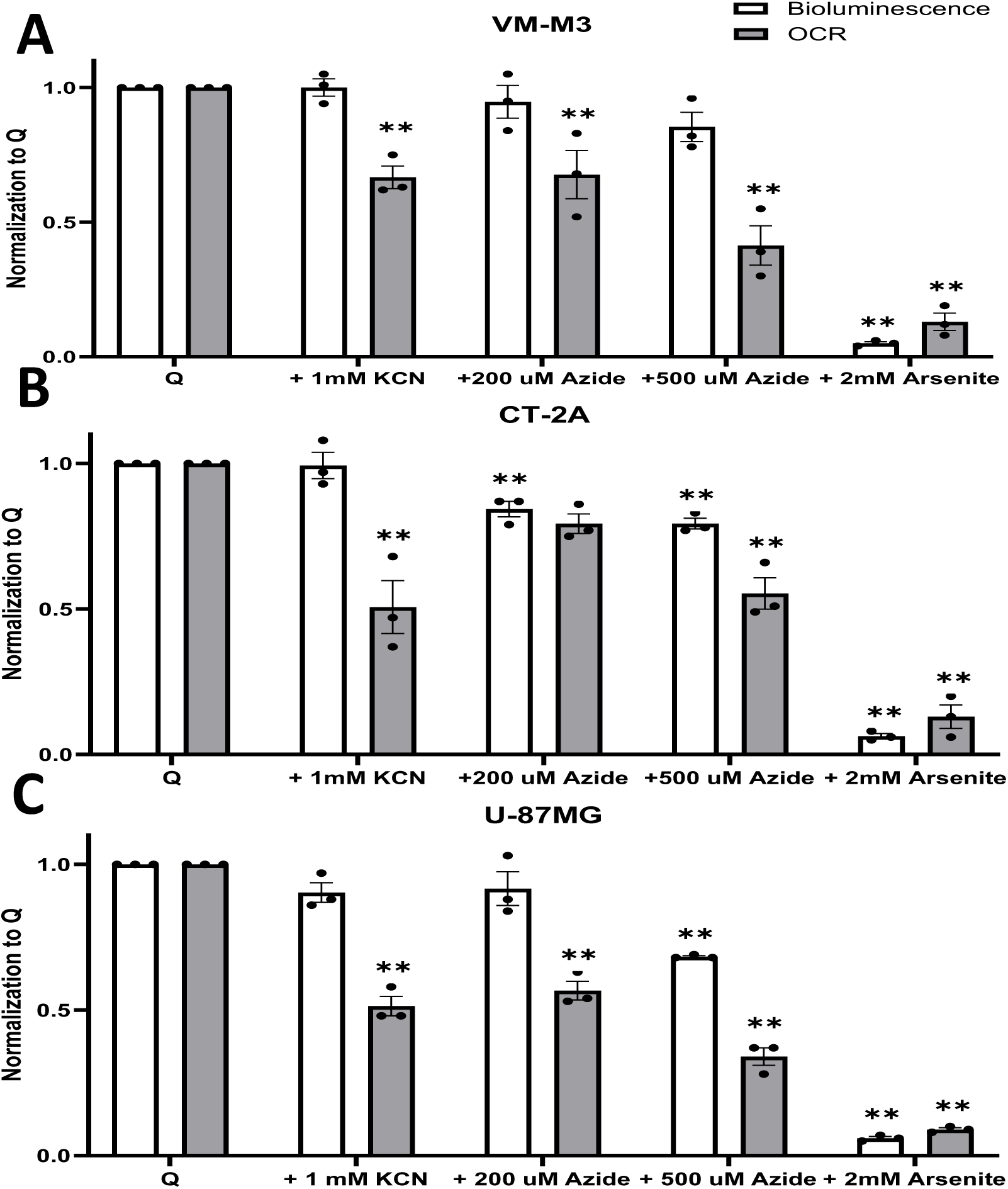
Oxygen Consumption is Independent of ATP Synthesis. A) VM-M3, B) CT-2A, and C) U-87MG cells cultured glutamine only media with sodium azide (200 or 500 µM), potassium cyanide (1 mM), or sodium arsenite (2 mM) for 6 hours. Black and grey bars indicate bioluminescence and oxygen consumption rate, respectively. All values are normalized and compared to control (Q) in three independent experiments. Values are plotted as mean ± SEM. Dunnett’s multiple comparison tests were used to evaluate statistical significance. Statistical significance is represented as: *p < 0.05; ** p < 0.01.

OCR was reduced by 21% and 45% when the CT-2A cells were cultured in media containing glutamine and 200 µM or 500 µM azide, respectively (**Figure 4B**). The reduction in OCR was significant at 200 µM and 500 µM. Bioluminescence decreased non-significantly at 200 µM yet was significant at 500 µM. Although 500 µM azide reduced both bioluminescence and oxygen consumption rate, the relative reduction in oxygen consumption (− 45%) was twice that of the reduction in bioluminescence (− 21%) indicating that the reduction in OCR was not proportional to the reduction in bioluminescence in this cell line. Like the findings in the CT-2A cells, OCR decreased significantly across both azide concentrations in the U-87MG cells. Although the addition of 200 µM azide did not reduce bioluminescence, bioluminescence was reduced at 500 µM, indicating that the U-87MG cells may be more sensitive than the mouse cells to higher concentrations of azide treatment (**Figure 4C**). To further evaluate the linkage of OCR to ATP content, we treated the cells with potassium cyanide, a well-studied inhibitor of complex IV. We performed a complex IV activity assay to confirm KCN inhibition. Ferrocytochrome C rapidly oxidized in control wells; however, oxidation was inhibited by the addition of 1 mM KCN (**Figure S5**). Potassium cyanide treatment significantly reduced OCR in each tumor cell line without a corresponding reduction in bioluminescence (**Figures 4A-C**). These findings further emphasize that the reduction in OCR was not proportional to the reduction in ATP content in these cells showing that OCR is not an accurate measure of ATP content in these cells.

### Sodium Arsenite Abolishes Bioluminescence and OCR

To examine the effect of glutaminolysis inhibition on the ATP-generating succinyl-CoA ligase reaction in the TCA cycle, we used sodium arsenite to inhibit mitochondrial dehydrogenases including the upstream alpha-ketoglutarate dehydrogenase (Lenartowicz, 1990). Both bioluminescence and OCR were nearly abolished from incubation in 2.0 mM of sodium arsenite in VM-M3 (**Figure 4A**), CT-2A (**Figure 2B**), and U-87MG cells (**Figure 4C**). These findings indicate that bioluminescence was reduced more following inhibition of glutaminolysis than following inhibition of complex IV in these glioma cells.

### Extracellular succinate as an end product of the glutaminolysis pathway

Global quantitative metabolomics was conducted to identify proteins involved in glutamine catabolism. VM-M3 cells were cultured in C^13^-glutamine for 18 hours. Extracellular C^13^-labeled succinate was observed in the extracellular matrix of VM-M3 cells via the glutaminolysis pathway (**Figure 5A**). Next, VM-M3 cells were incubated in hypoxia to determine if the presence of extracellular succinate persisted regardless of oxygen availability. Cells cultured in glutamine alone exported similar amounts of succinate in normoxia (21% O_2_) and in hypoxia (0.1% O_2_; **Figure 5B**). This result was confirmed using a high-throughput enzymatic assay kit (**Figure 5C**). This method has previously been used to measure succinate in human plasma of diabetic cohorts (Ceperuelo-Mallafre et al., 2019; Astiarraga et al., 2020).

**Figure 5.**
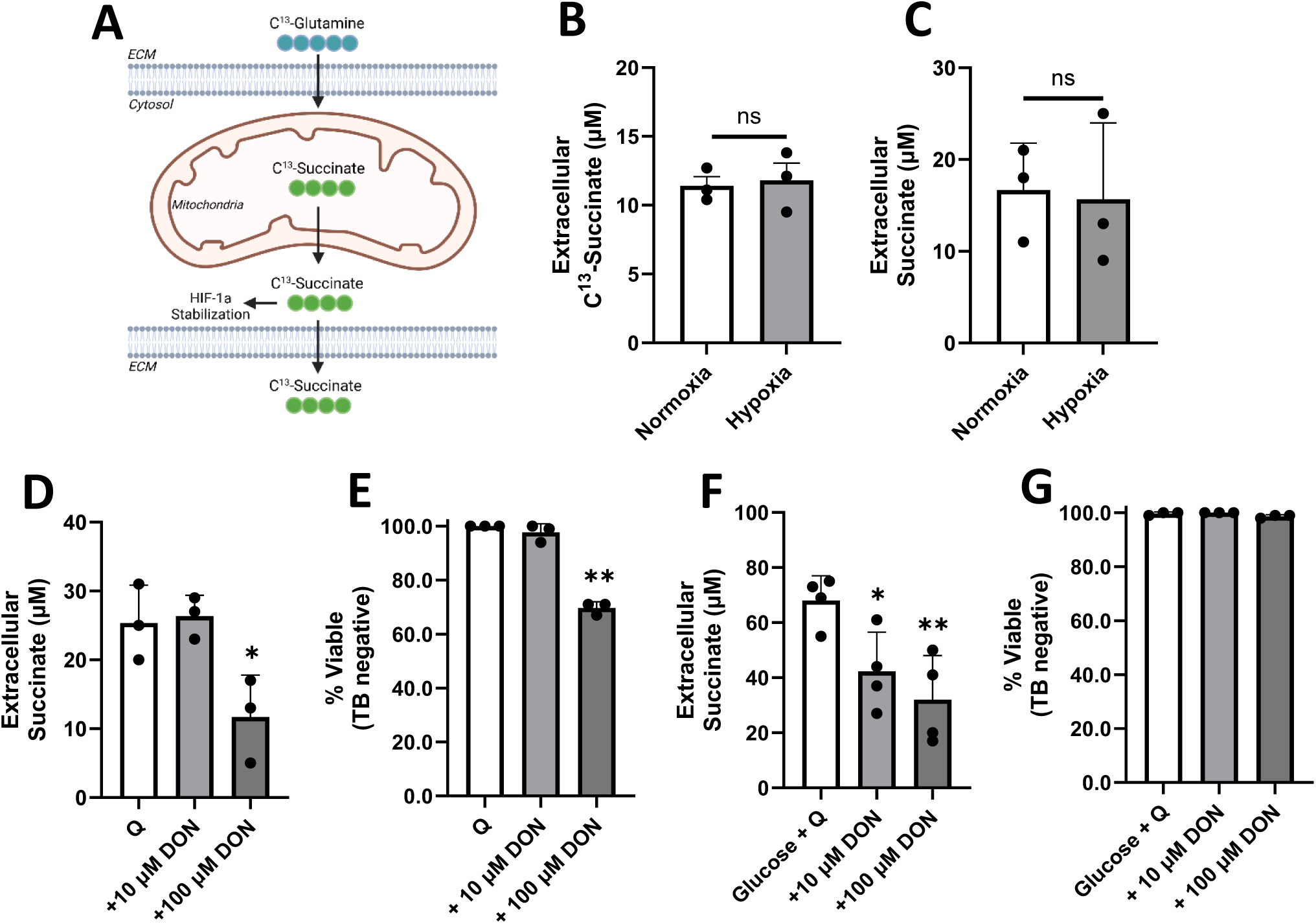
Labeled succinate as an end-product of glutamine breakdown. A) Simplified schematic of C^13^-glutamine tracing experiment. Glutamine enters the cell and is converted via glutaminolysis toward succinate. In GBM, succinate deviates from the TCA cycle and is exported from the mitochondria. Succinate has multiple fates within the cytosol including the stabilization of HIF-1a (Selak et al., 2005). Small amounts of remaining succinate are protonated and exported into the extracellular space. Image created using BioRender. B) Extracellular C^13^-succinate measured by mass spectrometry as described in *Materials and Methods*. C) Extracellular succinate measured by two-step colorimetric enzymatic assay. D) Extracellular succinate measured in the presence of 6-diazo-5-oxo-norleucine (DON) in glutamine only media after 6 hours. E) Trypan blue exclusion assay was performed on cells treated with DON in glutamine only media. F) Extracellular succinate measured in the presence of DON in glucose and glutamine containing media after 6 hours. G) Trypan blue exclusion assay was performed on cells treated with DON in glucose and glutamine containing media. Values are shown as mean ± SEM with three independent experiments. Unpaired student’s t-tests were used to evaluate statistical significance compared to control. Statistical significance is represented as: *p < 0.05; ** p < 0.01.

The pan-glutaminase inhibitor 6-diazo-5-oxo-norleucine (DON) was used to determine if succinate release could be reduced by blocking the first step in glutaminolysis. VM-M3 exposed to 100 µM DON significantly reduced the amount of extracellular succinate in the glutamine alone media (**Figure 5D**). Approximately 30% of VM-M3 cells became trypan blue positive, indicating that inhibition of glutaminolysis resulted in cell death (**Figure 5E**). This set of experiments was repeated in media containing both glucose and glutamine. Succinate release was markedly higher when glucose and glutamine were both available. This may be due to increased protonation of succinate which increases export of select compounds (Reddy et al., 2020; Murphy and Chouchani, 2022) or to a limited amount of pyruvate that is transported into the mitochondria (DeBerardinis et al., 2007). Succinate release was inhibited when exposed to 100 µM DON in the presence of glucose and glutamine (**Figure 5F**). However, trypan blue exclusion showed that nearly all cells remained viable (**Figure 5G**), suggesting that the availability of glucose preserved viability.

### Global metabolomics reveals multiple extracellular products of glutamine fermentation

Metabolomics analysis showed that the levels of several extracellular metabolites were greater in VM-M3 cells cultured in basal media containing glutamine than in basal media alone. (**Figure 6**). VM-M3 cells cultured in glutamine had significantly higher 2-hydroxyglutarate (2-HG) in the media than DMEM alone. 2-HG has been detected in previous studies, particularly in those bearing an *IDH1* mutation (Yan et al., 2009; Chesnelong et al., 2014). 2-HG is produced from the reductive pathway of the TCA cycle and has been shown to suppress HIF-1a activity (Bottcher et al., 2018). Extracellular asparagine was elevated from VM-M3 cells incubated in glutamine. This finding agrees with previous data showing that aspartate is coupled in a transamination reaction to form asparagine during glutamine catabolism (Panosyan et al., 2017). Similarly, alanine was increased which has been shown to be elevated during fermentative metabolism. Alanine accumulation results from transamination reactions during the breakdown of amino acids (Hochachka et al., 1975; Waterhouse et al., 1979; Tessem et al., 2008). This finding is consistent with a previous study that SF188 cells, cultured with [α-^15^N]glutamine, produced extracellular ^15^N-alanine (DeBerardinis et al., 2007). 2-methylglutaric acid was increased in samples containing glutamine. 2-methylglutaric acid is produced as a branchpoint from alpha-ketoglutarate (Goodman et al., 1975), but has not been thoroughly studied. Glutaric acid was observed in the urine of patients with cervical cancer (Liang et al., 2014). The precise characterization of glutaric acid remains to be elucidated, however its close relationship to the glutaminolysis pathway poses an intriguing avenue for further research.

**Figure 6.**
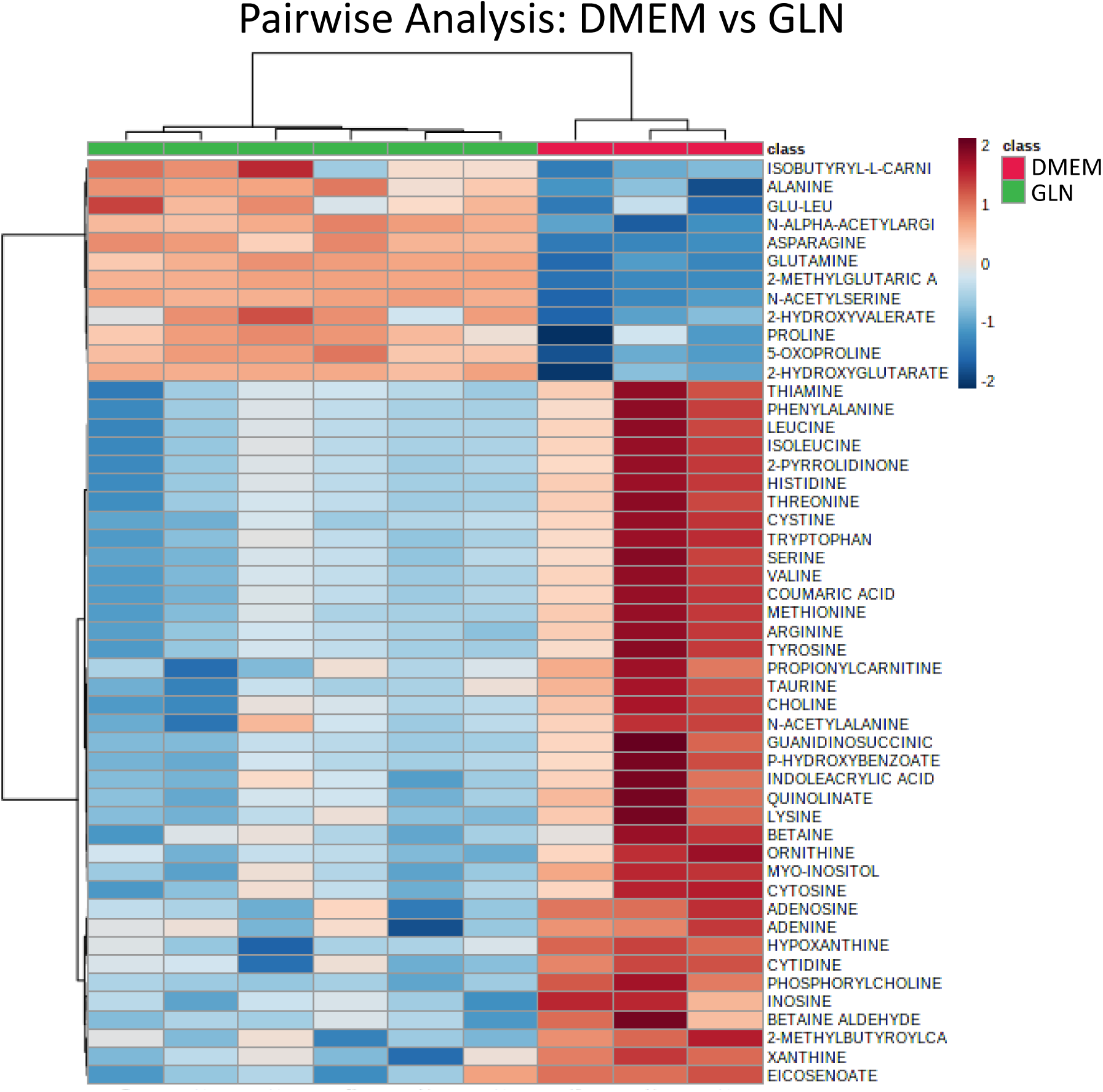
Clustering analysis of differentially expressed metabolites by presence or absence of glutamine in VM-M3 cells. A) Metabolites quantified using mass spectrometry and analyzed by MetaboAnalyst 4.0 to generate a heat map ranked by fold change. Each column represents one sample with rows associated with the indicated metabolite. Red represents the fold change increase while blue represents the fold change decrease based on the presence (green, GLN) or absence (red, DMEM) of glutamine.

The global metabolomics approach was extended to five conditions (**Figure 7A**). Three of the conditions were basal media, glutamine alone, or glucose and glutamine in normoxia (21% O_2_). The remaining two conditions were glutamine alone or glucose plus glutamine in hypoxia (0.1% O_2_). Lactate production was significantly greater under conditions with glucose than under conditions in the absence of glucose (glutamine alone) in support of our previous findings (Ta and Seyfried, 2015; Seyfried et al., 2020). All conditions showed a significant increase in N-acetylserine relative to basal media. N-acetylserine is commonly observed in cancers and has been proposed as a biomarker in endometrial carcinoma (Shao et al., 2016). Glucosamine, an amino sugar responsible for glycosylation, was present in our samples containing glucose and glutamine regardless of oxygen availability. Glycosylation can upregulate glycolytic enzymes (Lam et al., 2021) potentially to support the compensatory fermentation observed in malignant glioma cells. The nucleosides cytidine, uracil, and adenosine were elevated in samples containing glutamine exposed to hypoxia suggesting differential partitioning of glutamine catabolism depending on oxygen availability. Adenosine has been implicated as a key signaling molecule in response to metabolic stress in cancer cells (Yegutkin and Boison, 2022). Together, the metabolites that are abundant in the presence of fermentable fuels support a myriad of processes necessary for unbridled cellular division.

**Figure 7.**
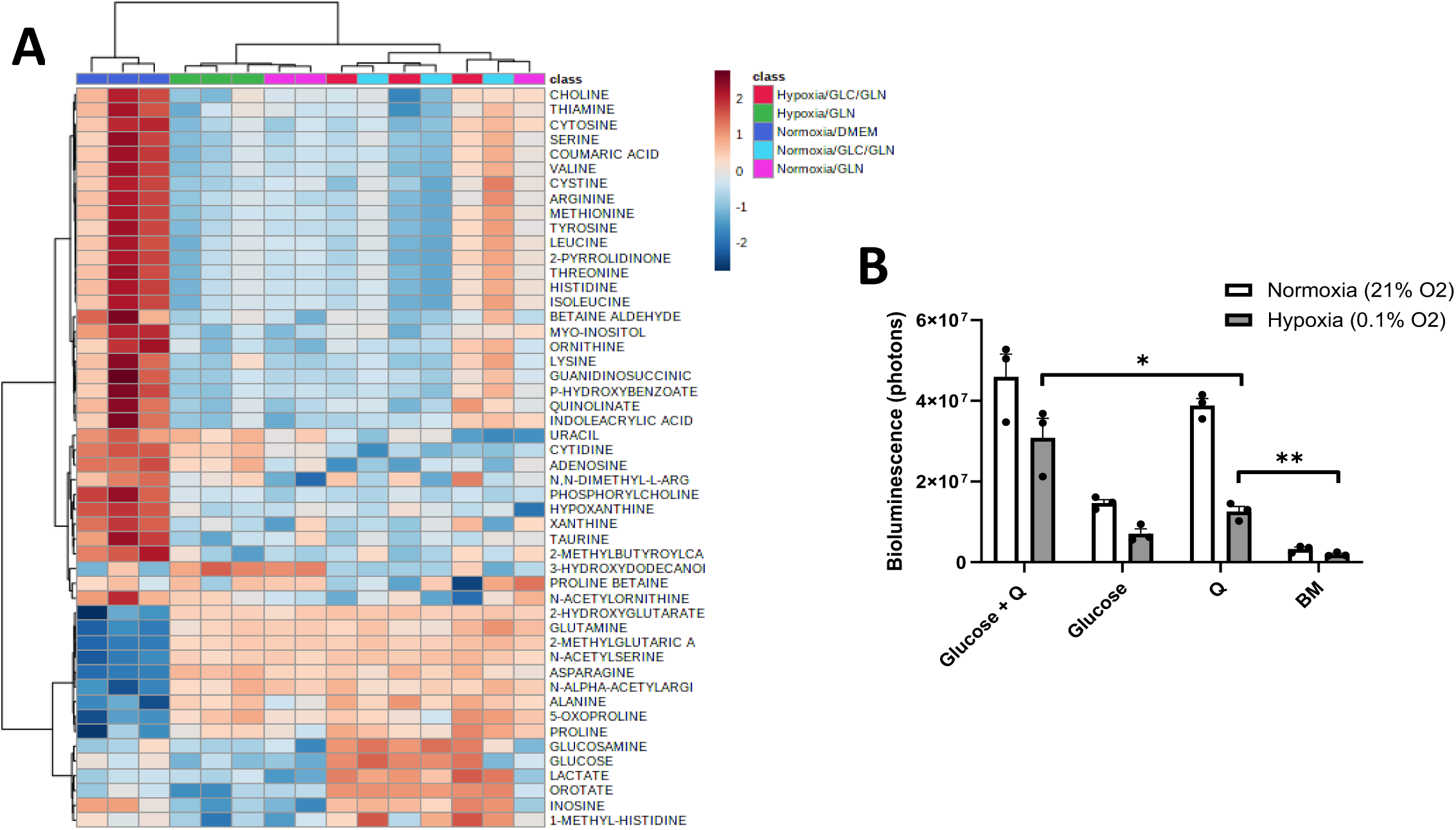
Global metabolomics of VM-M3 cells in hypoxia and normoxia. A) Metabolites were quantified using mass spectrometry and analyzed by MetaboAnalyst 4.0 as previously described. VM-M3 cells were grown in 12 mM glucose (GLC), 2 mM glutamine (GLN), or both. Cells were incubated in either normoxia (21% O_2_) or hypoxia (0.1% O_2_) for 18 hours. B) Bioluminescence measurements in VM-M3 cells cultured in hypoxia or normoxia after 18 hours. Values are shown as mean ± SEM with three independent experiments. Student’s t-tests were used to evaluate statistical significance: *p < 0.05; ** p < 0.01

### VM-M3 cells can sustain ATP content under hypoxia and glucose deprivation

It is well documented that cancer cells, including VM-M3 cells, can survive and proliferate for extended periods in hypoxic environments that contain glucose (Ta and Seyfried, 2015). We next evaluated the influence of deep hypoxia (0.1% O_2_) on ATP content and viability in VM-M3 cells cultured for 18 hours in the presence or absence of glucose and glutamine (**Figure 7B**). Bioluminescence was sustained in VM-M3 cells under severe hypoxia in the presence of glucose and glutamine. Cells grown in glutamine alone media maintained significantly higher bioluminescence than in the cells cultured in basal media alone. These data show that glutamine alone can maintain limited ATP content under severe oxygen deprivation. Viewed collectively, these findings present strong evidence showing that glutamine can support ATP production in mouse and human glioma cells through mitochondrial substrate level phosphorylation in the glutaminolysis pathway.

## Discussion

The objective of this research was to evaluate the origin of energy production in mouse and human glioma cells. Without energy (ATP), no cell can survive or grow making energy production a central issue in glioma cell viability and growth. The origin of ATP in cancer has been controversial since Warburg first proposed that cancer cells synthesize energy primarily through fermentation following irreversible damage to mitochondrial oxidative respiration (Burk and Schade, 1956; Warburg, 1956; Weinhouse, 1956; Aisenberg, 1961; Koppenol et al., 2011; Seyfried et al., 2020). In this study, we present evidence showing that glucose and glutamine together are necessary and sufficient for maintaining ATP content through fermentation in three glioma cell lines that differed in cell biology, genetic background, and species of origin. Although previous studies have viewed glutamine as a purely anaplerotic metabolite for maintaining ATP content through OxPhos in glioma (Wise and Thompson, 2010; Martins et al., 2020; Khadka et al., 2021), our data show that glutamine could also maintain significant ATP content through mitochondrial substrate-level phosphorylation (mSLP) in the glutaminolysis pathway with succinate release into the microenvironment as an end-product. We found that glutamine, like glucose, could be fermented for maintaining ATP content in either the presence or absence of oxygen. ATP content persisted for up to 24 hours in the tumor cells in serum-free media (either DMEM or PBS) containing glutamine alone, thus eliminating compensatory ATP from glucose-driven glycolysis (Warburg effect). Sufficient ATP content was also observed in glioma cells grown under deep hypoxia (0.1% oxygen) and in the absence of glucose thus excluding both OxPhos and glycolysis as significant sources of ATP under these conditions. Previous studies showed that tumor cells, including gliomas, could survive in the presence of electron transport chain inhibitors, e.g., rotenone, cyanide, and oligomycin, as long as fermentable fuels were also present in the growth environment (Warburg, 1927; Barron and Hoffman, 1930; Warburg, 1931; Ceruti et al., 2005; Hao et al., 2010; Renner et al., 2010; Ta and Seyfried, 2015). Our data support these findings and show for the first time that glutamine could be fermented for sufficient ATP content in glioma cells. Also consistent with recent findings in non-neural tumors (Doczi et al., 2023), our data show that ATP contribution through OxPhos was neither necessary nor sufficient for the growth of the VM-M3, CT-2A, or U-87MG glioma cells.

Amino acids have become prevalent targets for therapeutic approaches that aim to exploit vulnerabilities in cancer metabolism (Bhutia et al., 2015). To determine if any non-essential or essential amino acid could match the effect of glutamine for maintaining ATP content in glioma cells, we evaluated the effect of the other 19 amino acids used either alone or in combination with glucose on ATP-dependent bioluminescence. Glioma cells were grown either in DMEM or in PBS without serum to exclude any other macro- or micro-nutrients that might confound data interpretation. The results showed that ATP content and survival was greater for glutamine and glutamate than for any of the other 18 amino acids independent of glucose availability in the media. It is important to mention that glutamine and glutamate are the only amino acids that can maintain net ATP content through the glutaminolysis pathway without involving energy expenditure for entry into the TCA cycle (Seyfried, 2010; Chinopoulos and Seyfried, 2018). A greater number of glutamine transporters than glutamate transporters might account in part for the greater effect of glutamine over glutamate for maintaining ATP content in these glioma cells (Bhutia et al., 2015). While none of the other amino acids could maintain ATP content to a degree equivalent to glutamine or glutamate, we found that high concentrations of the sulfur-containing amino acids, methionine and cysteine, inhibited ATP content in the glioma cells tested. Hence, glutamine had the greatest effect in providing sufficient ATP content in these glioma cells.

To determine if oxygen consumption rate (OCR) was an accurate measurement for ATP content through OxPhos in the three glioma cell lines, we used a novel experimental design in measuring OCR in tandem with ATP-dependent bioluminescence over 6 hours. The data showed that sodium azide and potassium cyanide significantly lowered OCR without significantly reducing bioluminescence in the presence of glutamine alone. While OCR might be a good marker for ATP content in normal respiring tissues (Tolentino et al., 2006; Cadenas, 2018), our findings showed that OCR was not an accurate marker for ATP content through OxPhos in the three cell lines that we evaluated. Our findings agree with those of other studies indicating that many tumor cells might use oxygen for activities other than ATP maintenance, e.g., ROS production (Piccoli et al., 2005; Ramanathan et al., 2005; Herst and Berridge, 2007; Zhou et al., 2011). Additionally, the proton motive gradient can be dissipated through uncoupling mechanisms resulting in a mismatch of oxygen consumption and complex V-linked proton translocation (Arcos et al., 1969; Hall et al., 2013; Leznev et al., 2013; Velez et al., 2013; Vozza et al., 2014; Pacini and Borziani, 2016; Zhao et al., 2019). Caution is therefore necessary in considering OCR as an accurate biomarker for ATP content through OxPhos in cultured glioma cells especially if fermentable fuels like glucose and glutamine are also present in the media.

We previously suggested that ATP could be synthesized in glioblastoma through mitochondrial substrate-level phosphorylation (mSLP) in the glutamine-driven glutaminolysis pathway (Chinopoulos and Seyfried, 2018; Seyfried et al., 2020). Previous studies found that oxygen-independent mSLP was a major source of ATP content necessary for the growth of the parasite *Trypanosoma brucei* (Bochud-Allemann and Schneider, 2002; Taleva et al., 2023). Additionally, mSLP could rescue proliferation in respiration-impaired yeast by maintaining the mitochondrial membrane potential (Schwimmer et al., 2005). Previous studies in non-neural cancer cells also provided evidence for a role of mSLP in driving tumor growth (Gao et al., 2016). We found that VM-M3 cells were more sensitive to inhibition of glutamine catabolism upstream of succinyl-CoA synthetase using the alpha-ketoglutarate dehydrogenase inhibitor sodium arsenite than from inhibition of complex IV using potassium cyanide. In addition to maintaining oxygen-independent ATP content for glioma cell viability, ATP production through glutamine-driven mSLP can also reduce ATP hydrolysis from a reversal of the ATP synthase reaction during bioenergetic stress (Chinopoulos et al., 2010; Doczi et al., 2023; Ravasz et al., 2024). Our results suggest that glutamine, which has been considered primarily an anaplerotic respiratory fuel for growth, can also be fermented via mSLP, as an energy mechanism that has been largely unrecognized in cancer, especially in gliomas (Wise and Thompson, 2010; Martins et al., 2020; Garcia et al., 2021; Khadka et al., 2021; Ravasz et al., 2024).

Succinate is the known end-product of the glutamine-driven glutaminolysis pathway (Chinopoulos, 2019, 2020). We and others showed that little lactate is produced from glutamine in tumor cells (Portais et al., 1996; Ta and Seyfried, 2015) relative to glucose. Our ^13^C labeling and DON inhibition experiments showed that the extracellular succinate produced in the VM-M3 cells was derived from the glutamine added to the culture media, thus supporting the origin of succinate through the glutaminolysis pathway in these cells. Additionally, succinate can be derived from malate due to complex II reversal in hypoxia (Ravasz et al., 2024). Succinate, like lactate, accumulates extracellularly following acute inhibition of OxPhos under conditions of prolonged diving (Hochachka et al., 1975), ischemia (Taegtmeyer, 1978; Zhang et al., 2018; Chinopoulos, 2019), hemorrhagic shock (Taghavi et al., 2022), and high-intensity muscle exercise (Reddy et al., 2020). The accumulation of these fermentation end products are biomarkers for compensatory ATP maintenance through substrate-level phosphorylation in the cytoplasm and the mitochondria, respectively. It is noteworthy that the accumulation of these biomarkers ceased following the resumption of OxPhos activity indicating their linkage to transient ATP maintenance through fermentation. In contrast to the transient accumulation of fermentation end products seen in oxygen deprived normal cells and tissues, the glioma cells we studied continued to produce lactate and succinate even in the presence of oxygen due largely to inefficient OxPhos.

We also found that both the PKM1 and PKM2 isoforms were expressed in VM-M3 cells as was previously demonstrated in the CT-2A and the U-87MG cells (Marsh et al., 2008; Gao et al., 2021). These findings are consistent with previous studies showing that the PKM2 isoform is expressed in various human glioma cell lines (Luan et al., 2015). Unlike the PKM1 isoform, which produces ATP in the glycolytic pathway, the dimerized PKM2 isoform exists as a low-affinity enzyme which diverts phosphoenolpyruvate toward auxiliary pathways that result in lactate production while circumventing the net ATP step in glycolysis (Liu and Vander Heiden, 2015; Chinopoulos, 2020; Seyfried et al., 2020; Liu et al., 2022). Hence, caution is necessary in considering lactate as a quantitative biomarker for ATP content through glycolysis in glioma cells that express the PKM2 isoform.

The lactate and succinate accumulation in tumor cells has been linked to the well-known abnormalities in the number, structure, and function of mitochondria, which would cause OxPhos inefficiency (Zhou et al., 2011; Chinopoulos, 2020; Seyfried et al., 2020). Abnormalities in the electron transport chain were previously found in the VM-M3, the CT-2A, and the U-87MG cells that would compromise the efficiency of OxPhos in these glioma cells (Kiebish et al., 2008; Zhou et al., 2011; Nelson et al., 2021). Besides serving as a biomarker for mitochondrial fermentation, succinate is also known to stabilize Hif-1a through inhibition of the prolyl hydroxylase complex (Selak et al., 2005). A new model is presented in **Figure 8** highlighting the synergistic interaction between the high-throughput glycolysis and glutaminolysis pathways that drive the dysregulated cell growth and maintain ATP content through the aerobic fermentation of both glucose and glutamine.

**Figure 8.**
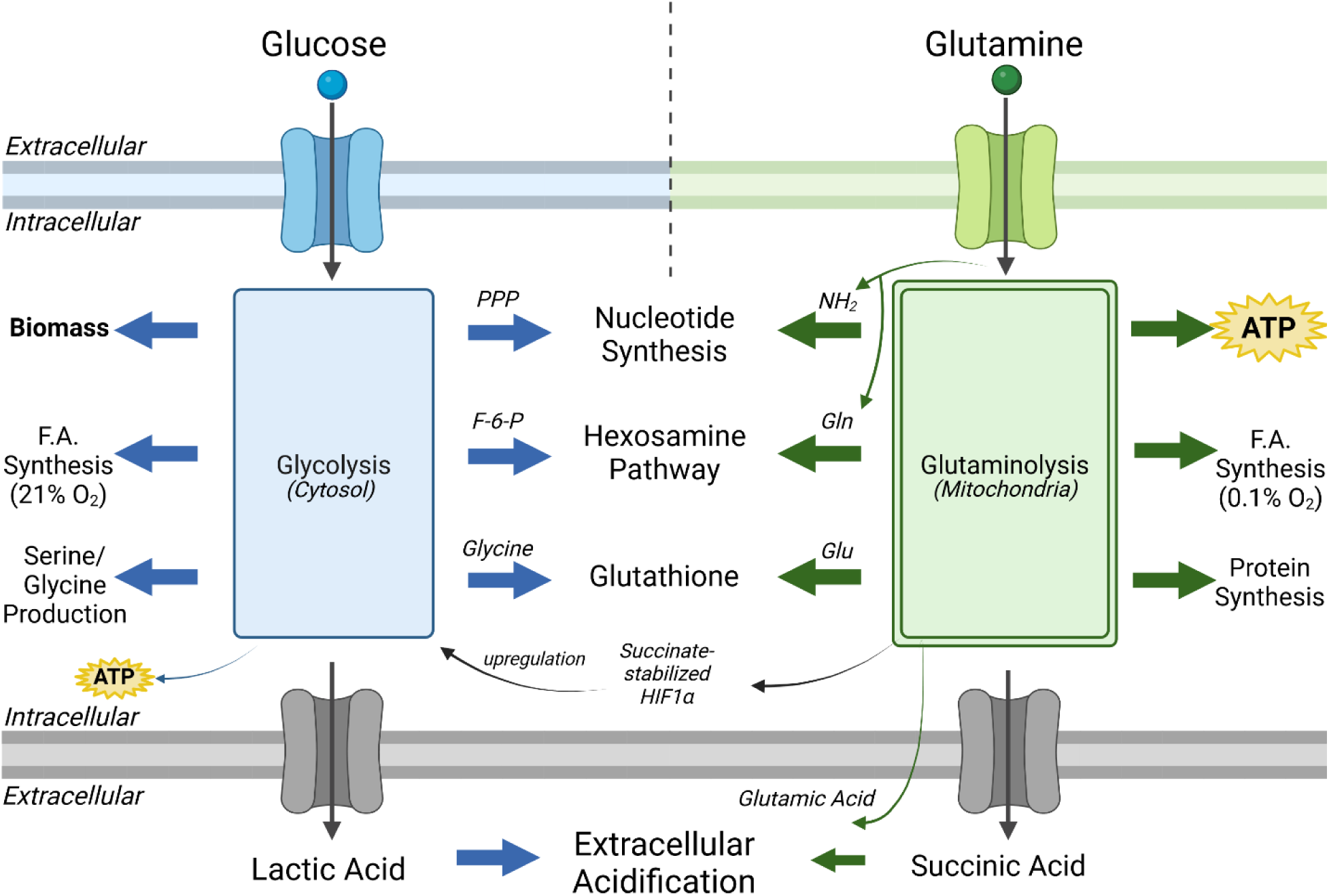
High-throughput synergy between the glycolysis and the glutaminolysis pathways drive the dysregulated growth of glioma cells. Glioma cells are dependent on both glucose and glutamine for maintaining dysregulated growth. This dual dependency can be explained largely through the synergy between these two pathways that facilitate the synthesis of ATP and biomass. Glucose (blue) enters the cell through GLUT1 transporters and is metabolized through the Embden-Meyerhof-Parnas glycolytic pathway. This 10-step pathway contributes to several pro-biomass pathways including the pentose phosphate pathway (PPP) for nucleotide synthesis, the hexosamine pathway for N- and O-linked glycosylation precursors, glycine synthesis for glutathione production, and serine for one-carbon metabolism. Some glucose carbons are diverted to synthesize fatty acids in normoxia. Glucose carbons that reach pyruvate kinase are exported as lactate which contributes to extracellular acidification. Glutamine (green) enters the cell primarily through the SLC1A5 transporter and enters the glutaminolysis pathway. Glutamine is essential for producing glucosamine-6-phosphate, a key intermediate in the hexosamine pathway that contributes to N- and O-linked glycosylation. The amide nitrogen released from the conversion of glutamine to glutamate contributes to nucleotide synthesis. Glutamate is combined with glycine and cysteine to form glutathione, an antioxidant that protects tumor cells from ROS. The remaining glutamate is converted first to alpha-ketoglutarate (a-KG). a-KG will divert in the reductive TCA cycle through citrate and be used for fatty acid synthesis in hypoxia. Otherwise, a-KG follows the oxidative pathway and is converted to succinyl-CoA. Succinyl-CoA is the substrate for mitochondrial substate level phosphorylation (mSLP) that produces ATP and succinate. Succinate has been shown to stabilize HIF1a via inhibition of prolyl hydroxylase (Selak et al., 2005) – a key protein that upregulates glycolysis. Lastly, both succinate and glutamate have been found to be excreted into the extracellular matrix and contribute toward acidification of the microenvironment. Figure created using BioRender.

## Conclusions

Our data show that glutamine could maintain sufficient ATP content in either the presence or absence of oxygen through mitochondrial substrate-level phosphorylation in the glutaminolysis pathway during glucose deprivation. No other amino acid had a stimulatory effect on ATP content comparable to that of glutamine in the mouse and human malignant glioma cells that we evaluated. The data show that oxygen consumption and lactic acid production are not accurate quantitative measures of ATP content through OxPhos or glycolysis, respectively, in these glioma cell lines. We also found that ATP content through OxPhos alone was unable to maintain bioluminescence of glioma cells in the absence of glucose and glutamine, indicating that these metabolites are both necessary and sufficient for maintaining growth of these glioma cells. A new metabolic model is presented highlighting the synergistic interaction between the high-throughput glycolysis and glutaminolysis pathways that converge on anabolic intermediates and maintain ATP content through the aerobic fermentation of both glucose and glutamine. Finally, the findings from this study can provide a general mechanism explaining the therapeutic efficacy seen from *in vivo* studies where the simultaneous restriction of glycolysis and glutaminolysis under nutritional ketosis enhance survival of young and adult mice bearing malignant brain tumors (Mukherjee et al., 2019; Seyfried et al., 2021; Seyfried et al., 2022; Mukherjee et al., 2023).

## Limitations

One limitation of our study involved the difficulty in determining the fraction of ATP that was synthesized within the mitochondria from OxPhos or mSLP given the metabolic entanglement of these two ATP sources. Consequently, we used a reductionist approach to evaluate ATP content through mSLP by growing tumor cells in the absence of glucose or by treating them with well-known mitochondrial inhibitors. Although the ETC inhibition that we used in our experimental design (6 hours) was longer than the < 2 hours inhibition times used in most other studies, extended ETC inhibition studies would be warranted to better evaluate the consequences of long-term ETC inhibition on tumor cell growth and viability. Finally, we limit our conclusions only to the mouse and human glioma cell lines that we studied. It is not clear if similar results would be obtained using our experimental design for other cancer cell lines. Comparison of nutrient utilization between syngeneic primary cells and cancer cells of similar cell origin may provide insight into exploitable bioenergetic targets of carcinogenesis.

## Author Contributions

DCL, LT, PM, TD, CC, and TNS contributed to the conceptualization and design of the research. DCL, LT, and PM performed the bulk of experiments and acquired the data. MD, BG, SK, NRN, and MK generated metabolite data sets. DCL and TNS drafted and revised the manuscript. All authors have read and agreed to the published version of the manuscript.

## Data Availability Statement

All data supporting the findings of this study are available from the corresponding author upon reasonable request.

## Acknowledgements

We thank the Foundation for Metabolic Cancer Therapies, CrossFit Inc., Dr. Joseph C. Maroon, Dr. Edward Miller, The Broken Science Initiative, Children with Cancer UK, The Corkin Family Foundation, and the Boston College Research Expense Fund for their support. We also thank Miguel Estevez for providing the U-87MG cell line. We thank Dr. Laura Shelton for her original background work on the subject. We thank Bret Judson and the Boston College Imaging Core for their assistance with microscopy. Lastly, we thank the three anonymous reviewers for their input and suggestions, which improved the quality of our manuscript.

