## Supplemental Figures and Tables for "Amino Acid and Glucose Fermentation Maintain ATP Content in Mouse and Human Malignant Glioma Cells"

#### Supplemental Figure 1

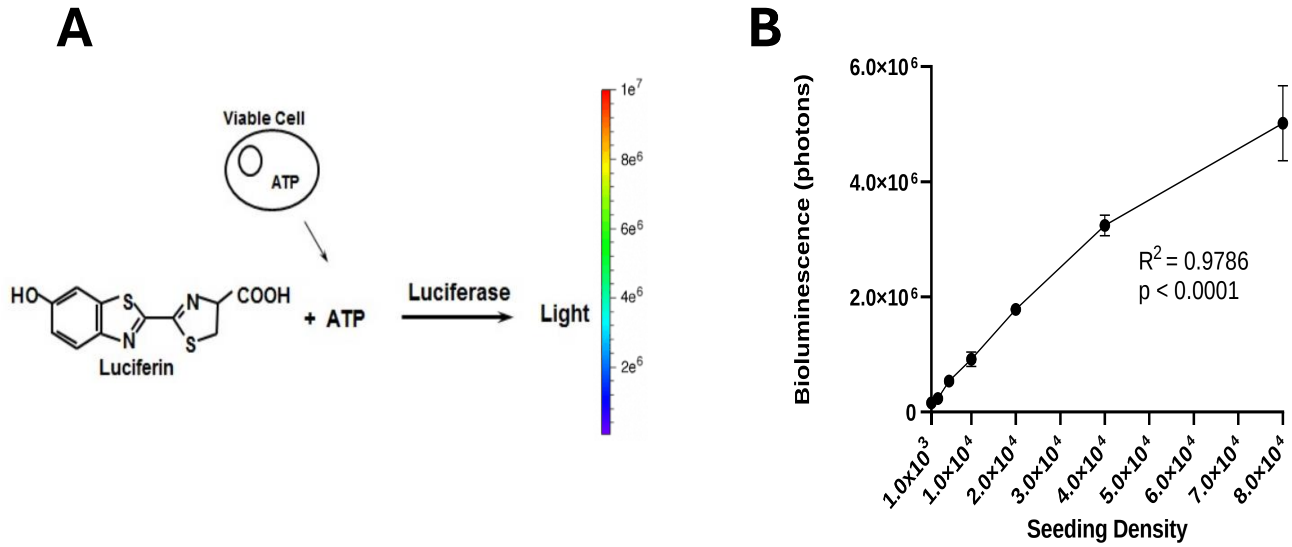


**Supplemental Figure 1. Correlation between seeding density and bioluminescence.** A) Schematic of ATP-dependent luciferin-luciferase bioluminescent reaction. Modified from *Cell Viability Assays*. B) VM-M3 cells were seeded in seeding media and adhered for 6 hours. Media was swapped from seeding media to experimental media (without phenol red and serum) immediately prior to bioluminescence reading. Luciferin (10 mg/mL) was applied, and the reading was taken after five minutes. 4-5 measurements in 96-well plates were used for each seeding density. Pearson's correlation analysis was performed with a 95% confidence interval (0.9261-0.9985).

#### Supplemental Figure 2

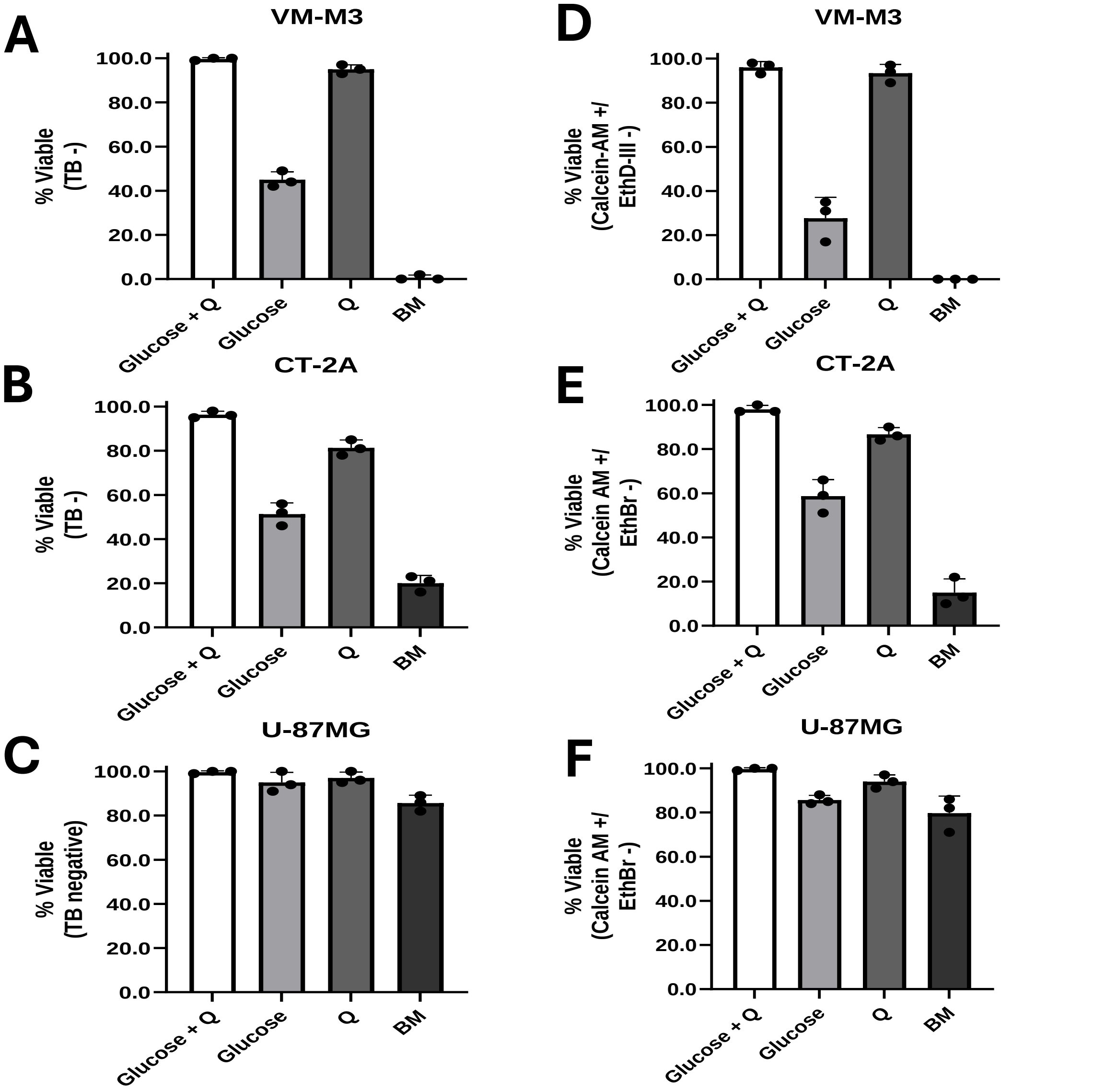


**Supplemental Figure 2. Influence of glucose and glutamine on cell viability.** A) VM-M3, B) CT-2A, and C) U-87MG cells were seeded at a density of 5.0 x $10^3$ and cultured for 24 hours with indicated media compositions. Glucose was added at 12 mM and glutamine (Q) was added at 2 mM. BM represents DMEM with no added glucose or glutamine. Trypan blue exclusion assay was used to determine viability. D) VM-M3, E) CT-2A, and F) U-87MG cells were cultured as described in A-C. Calcein-AM and ethidium homodimer III (EthD-III) staining assay was used to determine viability. Data are shown as mean ± SEM with three independent experiments.

#### Supplemental Figure 3

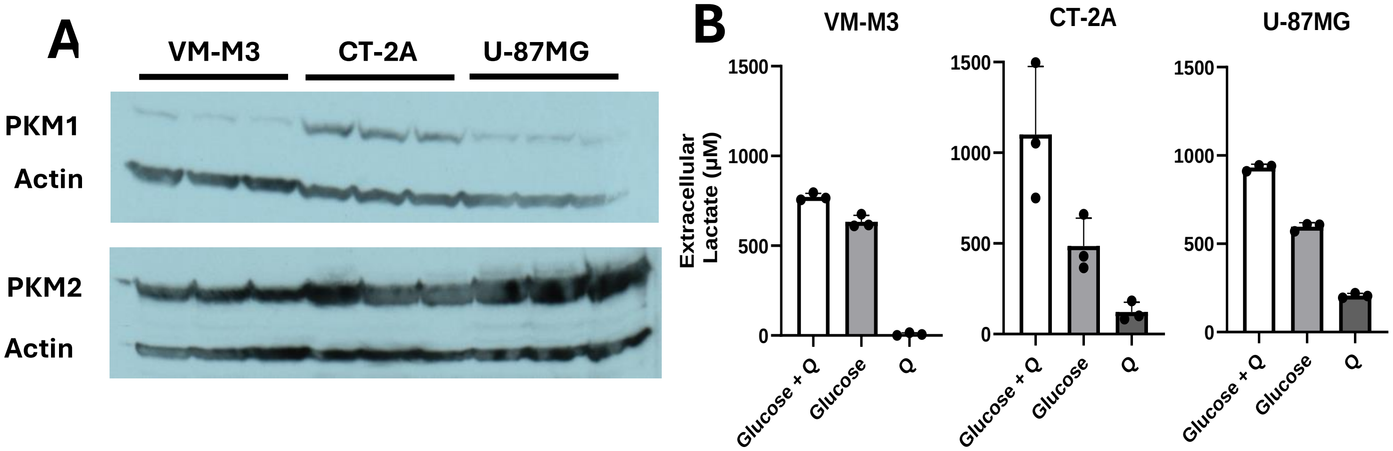


**Supplemental Figure 3. Presence of PKM2 in VM-M3, CT-2A, and U-87MG cells.** A) Representative western blots of PKM1 and PKM2 expression in VM-M3, CT-2A, and U-87MG cells. Actin was used as a loading control. B) Extracellular lactate produced by glucose, glutamine, or both after 6 hours. Lactate was measured by colorimetric enzymatic assay. Background reading (basal media) was subtracted from all values. Bars represent three independent experiments.

#### Supplemental Figure 4

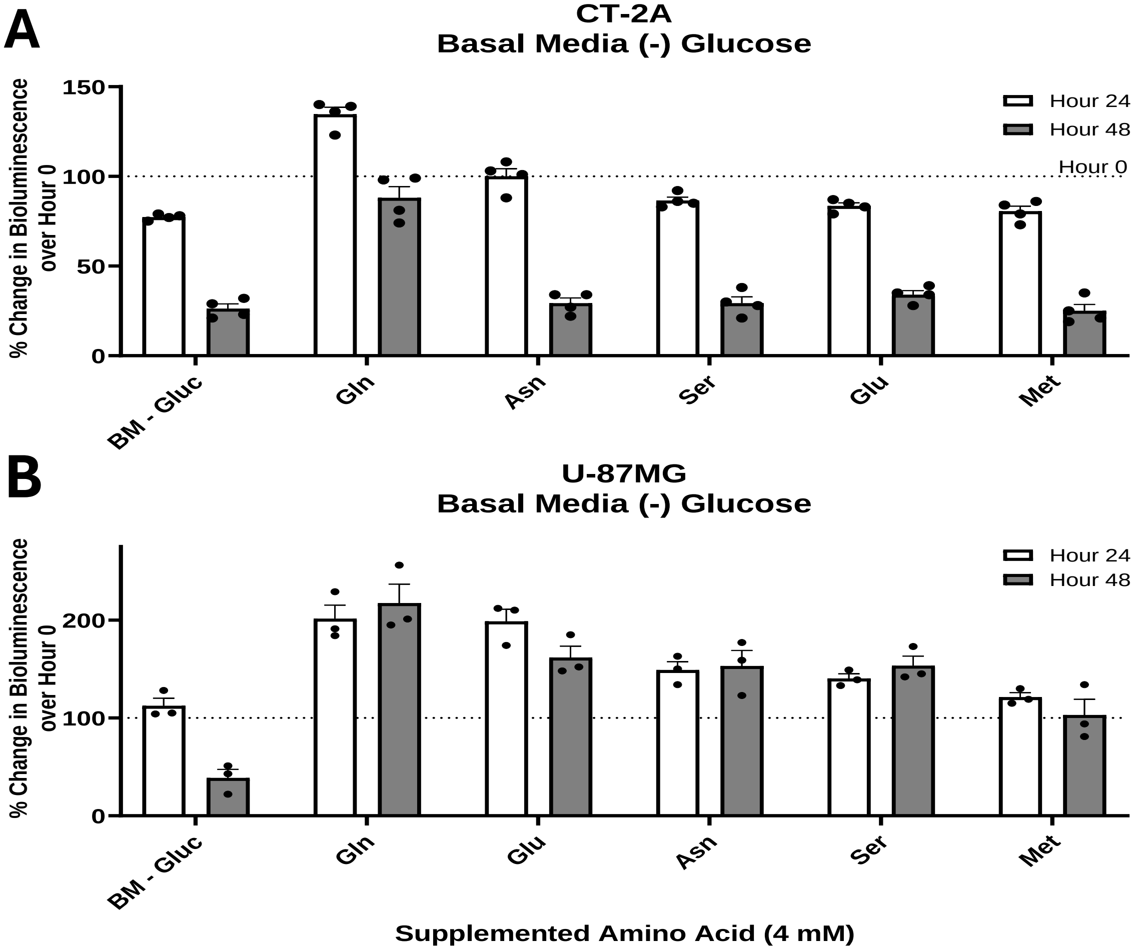


**Supplemental Figure 4. Influence of supplemented amino acids on bioluminescence in basal media.** A) CT-2A and B) U-87MG cells cultured in basal media with the indicated amino acids supplemented at 4 mM. Amino acids are ordered by descending percent change at hour 24. Cells were seeded at a density of 1.0 x $10^4$ cells/well. The bioluminescence value at hour 0 is represented by a dashed line. Values are measured as mean ± SEM with three or four independent experiments.

#### Supplemental Figure 5

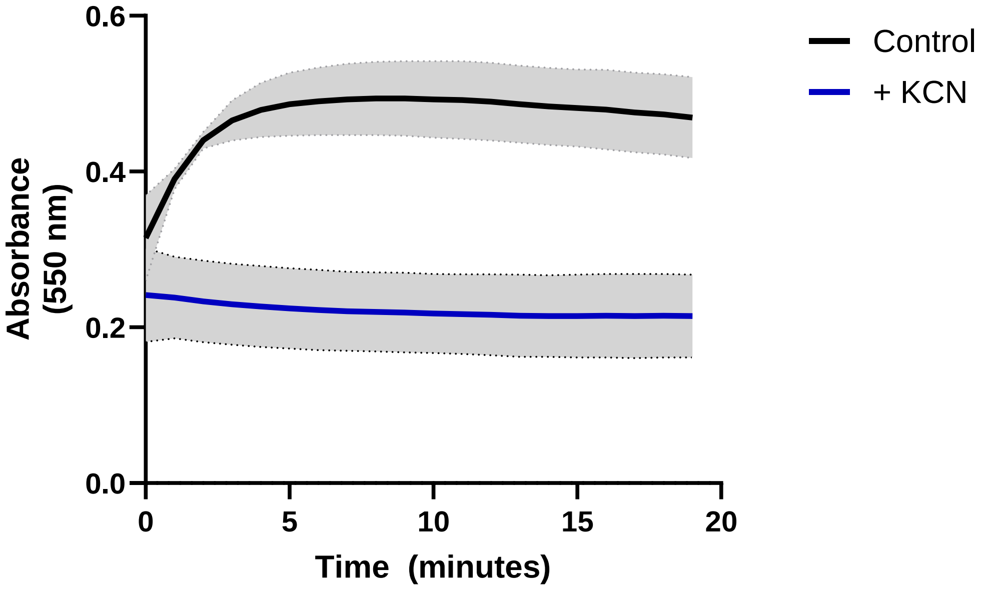


**Supplemental Figure 5. Influence of Potassium Cyanide on Complex IV Activity.** VM-M3 cells were cultured with (blue) or without (black) the addition of 1 mM KCN. The absorbance was measured at 550 nm for 20 minutes upon the addition of 50 µM ferrocytochrome C. Values are shown as mean ± SD with three independent experiments.

### Tables

| Amino Acid | 24 Hours | | 48 Hours | |
|---|---|---|---|---|
| | Photons[a] | % Change[b] | Photons[a] | % Change[b] |
| None† | 1.25e5 | -64 ± 3** | 2.06e4 | -94 ± 1 |
| Glutamine | 6.18e5 | +91 ± 4** | 3.19e5 | 0 ± 1** |
| Glutamic Acid | 4.11e5 | +30 ± 4** | 1.95e5 | -38 ± 4** |
| Lysine | 4.00e5 | +10 ± 6 ** | 1.23e5 | -66 ± 3** |
| Phenylalanine | 3.32e5 | -9 ± 5** | 3.86e4 | -89 ± 1 |
| Threonine | 2.94e5 | -19 ± 3** | 5.35e4 | -85 ± 2 |
| Asparagine | 2.37e5 | -23 ± 7** | 5.03e4 | -84 ± 0 |
| Histidine | 2.83e5 | -22 ± 4** | 9.34e4 | -74 ± 4** |
| Isoleucine | 2.68e5 | -38 ± 6** | 5.39e4 | -88 ± 2 |
| Tyrosine | 1.90e5 | -48 ± 2* | 2.48e4 | -93 ± 1 |
| Aspartic Acid | 1.45e5 | -49 ± 11 | 2.99e4 | -89 ± 2 |
| Glycine | 1.38e5 | -55 ± 3 | 1.31e4 | -96 ± 0 |
| Arginine | 1.13e5 | -63 ± 7 | 1.09e4 | -96 ± 0 |
| Tryptophan | 1.03e5 | -71 ± 1 | 3.51e4 | -90 ± 1 |
| Serine | 6.93e4 | -73 ± 4 | 4.99e3 | -98 ± 0 |
| Leucine | 8.55e4 | -74 ± 3 | 6.42e3 | -98 ± 0 |
| Alanine | 7.11e4 | -76 ± 2 | 1.70e4 | -94 ± 0 |
| Cysteine | 8.43e4 | -77 ± 2 | 1.07e4 | -97 ± 0 |
| Proline | 6.37e4 | -78 ± 3 | 5.21e3 | -98 ± 0 |
| Valine | 3.13e4 | -88 ± 2** | 2.29e3 | -99 ± 0 |
| Methionine | 4.26e4 | -89 ± 2** | 2.84e3 | -99 ± 0 |

**Table 1 –** All conditions contain only the indicated amino acid added at 4 mM to DMEM. Change to experimental media from seeding media is considered hour 0. Starting photon values range from 3.06e5 to 4.57e5 depending on experiment lot. [a] represents the mean bioluminescence photons. [b] represents the percent change in mean bioluminescence (± SEM) from the starting point (hour 0). e5 is equivalent to $10^5$. †Two-way ANOVA followed by post-hoc Sidak's comparison for statistical significance, $*p < 0.05$; $**p < 0.01$.

| Amino Acid | 24 Hours | | 48 Hours | |
|---|---|---|---|---|
| | Photons[a] | % Change[b] | Photons[a] | % Change[b] |
| None† | 1.07e5 | -78 ± 2 | 2.22e4 | -95 ± 1 |
| Glutamine | 7.37e5 | +55 ± 9 ** | 5.14e5 | +8 ± 10 ** |
| Glutamic Acid | 6.75e5 | +42 ± 9 ** | 4.35e5 | -9 ± 12 ** |
| Glycine | 5.09e5 | +7 ± 7 ** | 5.37e4 | -89 ± 1 |
| Asparagine | 4.70e5 | -1 ± 6 ** | 1.55e5 | -67 ± 3 ** |
| Serine | 4.25e5 | -11 ± 4 ** | 5.80e4 | -88 ± 2 |
| Proline | 3.35e5 | -30 ± 5 ** | 8.99e4 | -81 ± 5 |
| Aspartic Acid | 2.99e5 | -37 ± 5 | 1.05e5 | -78 ± 3 * |
| Threonine | 1.26e5 | -67 ± 3 | 7.14e4 | -82 ± 2 |
| Isoleucine | 1.20e5 | -75 ± 2 | 2.48e4 | -95 ± 0 |
| Leucine | 6.55e4 | -77 ± 1 | 5.71e4 | -85 ± 1 |
| Lysine | 6.51e4 | -78 ± 1 | 5.69e4 | -85 ± 0 |
| Alanine | 7.19e4 | -85 ± 1 | 7.85e3 | -98 ± 0 |
| Histidine | 5.40e4 | -89 ± 1 | 1.11e4 | -98 ± 0 |
| Valine | 4.94e4 | -90 ± 1 | 1.59e4 | -97 ± 0 |
| Arginine | 4.40e4 | -91 ± 0 | 8.62e3 | -98 ± 0 |
| Tyrosine | 3.11e4 | -92 ± 0 | 1.78e4 | -95 ± 0 |
| Phenylalanine | 2.49e4 | -94 ± 0 | 1.31e4 | -96 ± 0 |
| Methionine | 1.40e4 | -96 ± 0 * | 1.16e4 | -97 ± 0 |
| Tryptophan | 9.28e3 | -97 ± 0 ** | 6.13e3 | -98 ± 0 |
| Cysteine | 6.93e3 | -98 ± 0 ** | 4.26e3 | -98 ± 0 |

**Table 2** – All conditions contain only the indicated amino acid added at 4 mM added to PBS. Change to experimental media from seeding media is considered hour 0. Starting photon values range from 2.25e5 to 3.53e6 depending on experiment lot. [a] represents the mean bioluminescence photons. [b] represents the percent change in mean bioluminescence (± SEM) from the starting point (hour 0). e5 is equivalent to $10^5$. †Two-way ANOVA followed by post-hoc Sidak's comparison for statistical significance, *$p < 0.05$; **$p < 0.01$.

| Amino Acid | 24 Hours | | 48 Hours | |
| --- | --- | --- | --- | --- |
| | Photons[a] | % Change[b] | Photons[a] | % Change[b] |
| None† | 1.21e6 | -42 ± 3 | 8.48e4 | -61 ± 4 |
| Glutamine | 3.92e6 | +119 ± 8** | 7.80e5 | +192 ± 17** |
| Lysine | 4.44e5 | +64 ± 11** | 1.98e5 | -29 ± 7** |
| Isoleucine | 4.62e5 | +13 ± 4** | 1.17e5 | -71 ± 2 |
| Glutamic Acid | 1.82e6 | +4 ± 2** | 1.96e5 | -47 ± 2 |
| Histidine | 2.45e5 | -10 ± 5** | 1.08e5 | -60 ± 4 |
| Glycine | 1.72e6 | -12 ± 7** | 7.42e4 | -64 ± 4 |
| Threonine | 1.95e5 | -22 ± 4 | 1.06e5 | -60 ± 3 |
| Aspartic Acid | 1.22e6 | -31 ± 9 | 1.12e5 | -61 ± 2 |
| Phenylalanine | 1.83e5 | -32 ± 2 | 5.96e5 | -76 ± 2 |
| Asparagine | 1.50e6 | -33 ± 2 | 1.07e5 | -56 ± 3 |
| Serine | 1.52e6 | -37 ± 2 | 4.50e4 | -59 ± 21 |
| Proline | 1.44e6 | -42 ± 3 | 2.29e4 | -81 ± 2 |
| Leucine | 1.26e6 | -43 ± 3 | 3.44e4 | -78 ± 3 |
| Valine | 1.10e6 | -46 ± 2 | 3.25e4 | -80 ± 2 |
| Alanine | 1.27e6 | -50 ± 1 | 4.75e4 | -72 ± 3 |
| Arginine | 1.25e6 | -51 ± 2 | 4.77e4 | -76 ± 2 |
| Methionine | 1.65e5 | -63 ± 1 | 2.27e4 | -95 ± 0** |
| Tryptophan | 9.15e4 | -66 ± 2* | 4.46e4 | -80 ± 3 |
| Tyrosine | 8.47e4 | -69 ± 1** | 2.01e4 | -91 ± 1** |
| Cysteine | 6.52e4 | -76 ± 1** | 1.87e4 | -92 ± 1** |

**Table 3** – All conditions contain the indicated amino acid added at 4 mM and 12 mM glucose added to DMEM. Change to experimental media from seeding media is considered hour 0. Starting photon values range from 4.60e5 to 5.04e5 depending on experiment lot. [a] represents the mean bioluminescence photons. [b] represents the percent change in mean bioluminescence (± SEM) from the starting point (hour 0). e5 is equivalent to $10^5$. †Two-way ANOVA followed by post-hoc Sidak's comparison for statistical significance, *$p < 0.05$; **$p < 0.01$.

| Amino Acid | 24 Hours | | 48 Hours | |
|---|---|---|---|---|
| | Photons[a] | % Change[b] | Photons[a] | % Change[b] |
| None† | 3.53e4 | -86 ± 1 | 7.23e3 | 3 ± 0 |
| Glutamine | 5.23e5 | +103 ± 12** | 4.27e5 | +66 ± 11** |
| Glutamic Acid | 3.72e5 | +45 ± 12** | 2.27e5 | -12 ± 15** |
| Serine | 6.52e5 | +42 ± 5** | 3.92e5 | -15 ± 3** |
| Isoleucine | 6.46e5 | +41 ± 9** | 3.50e5 | -24 ± 6** |
| Asparagine | 3.47e5 | +35 ± 6** | 2.33e5 | -9 ± 8** |
| Glycine | 3.38e5 | +31 ± 12** | 1.79e5 | -30 ± 13** |
| Threonine | 3.83e5 | -17 ± 4** | 2.50e5 | -46 ± 5** |
| Alanine | 2.03e5 | -21 ± 8** | 1.77e5 | -31 ± 11** |
| Leucine | 3.51e5 | -24 ± 5** | 3.04e5 | -44 ± 7** |
| Aspartic Acid | 1.76e5 | -32 ± 5** | 4.35e4 | -83 ± 3 |
| Tyrosine | 2.30e5 | -50 ± 2** | 5.76e4 | -87 ± 1 |
| Histidine | 1.83e5 | -60 ± 3* | 3.72e4 | -92 ± 0 |
| Valine | 1.71e5 | -67 ± 3 | 8.02e4 | -83 ± 1 |
| Proline | 8.49e4 | -67 ± 2 | 5.44e4 | -79 ± 4 |
| Cysteine | 3.31e4 | -87 ± 1 | 1.45e4 | -94 ± 1 |
| Methionine | 3.24e4 | -87 ± 1 | 1.40e4 | -95 ± 1 |
| Phenylalanine | 4.63e4 | -90 ± 1 | 1.16e4 | -97 ± 0 |
| Lysine | 1.76e4 | -93 ± 0 | 1.16e4 | -95 ± 0 |
| Tryptophan | 2.33e4 | -95 ± 0 | 1.39e4 | -97 ± 0 |
| Arginine | 1.11e4 | -96 ± 0 | 8.39e3 | -97 ± 0 |

**Table 4** – All conditions contain only the indicated amino acid added at 4 mM and 12 mM glucose added to PBS. Change to experimental media from seeding media is considered hour 0. Starting photon values range from 4.60e5 to 5.04e5 depending on experiment lot. [a] represents the mean bioluminescence photons. [b] represents the percent change in mean bioluminescence (± SEM) from the starting point (hour 0). e5 is equivalent to $10^5$. †Two-way ANOVA followed by post-hoc Sidak's comparison for statistical significance, *$p < 0.05$; **$p < 0.01$.

#### Supplemental Table:

| Amino Acid | Source (Catalog) |
|---|---|
| Alanine | Thermo Fisher Scientific (56-41-7) |
| Arginine | Thermo Fisher Scientific (74-79-3) |
| Asparagine | Alfa Aesar (A15012) |
| Aspartic Acid | Sigma-Aldrich (A6683) |
| Cysteine | Alfa Aesar (J63745.14) |
| Glutamic Acid | Sigma-Aldrich (49621) |
| Glutamine | Sigma-Aldrich (G8540) |
| Glycine | Sigma-Aldrich (410225) |
| Histidine | Sigma-Aldrich (H-8125) |
| Isoleucine | Sigma-Aldrich (I2752) |
| Leucine | Alfa Aesar (J62824) |
| Lysine | Alfa Aesar (A16249) |
| Methionine | Sigma-Aldrich (M9625) |
| Phenylalanine | Alfa Aesar (A12328) |
| Proline | Thermo Fisher Scientific (A10199) |
| Serine | Alfa Aesar (J62187) |
| Threonine | Acros Organics (72-19-5) |
| Tryptophan | Thermo Fisher Scientific (J62508) |
| Tyrosine | Alfa Aesar (J63511) |
| Valine | Alfa Aesar (J62943) |

**Supplemental Table 1** – Amino acids and their sources used in this study.
